# Flash properties of Gaussia Luciferase are the result of covalent inhibition after a limited number of cycles

**DOI:** 10.1101/2020.10.01.322248

**Authors:** Fenne Marjolein Dijkema, Matilde Knapkøien Nordentoft, Anders Krøll Didriksen, Anders Sværke Corneliussen, Martin Willemoës, Jakob R. Winther

## Abstract

Luciferases are widely used as reporters for gene expression and for sensitive detection systems. While luciferases from firefly and *Renilla* have long been used for analysis of intracellular expression, the luciferase (GLuc) from the marine copepod *Gaussia princeps*, has gained popularity, primarily because it is secreted and displays a very high light intensity. Firefly luciferase is characterized by kinetic behavior which is consistent with conventional steady-state Michaelis-Menten kinetics (termed “glow” kinetics). GLuc, conversely, displays what has been termed “flash” kinetics which signify a burst in light emission followed by a rapid decay. As the mechanistic background for this behavior is poorly characterized, we decided to decipher the mechanism in more detail. We show that decay in light signal is not due to depletion of substrate, but rather is caused by the irreversible inactivation of the enzyme. Inactivation takes place after between 10 and 200 reaction cycles, depending on substrate concentration. We found that the rate of inactivation is described by the sum of two exponentials with associated rate constants. The dominant of these of these increases linearly with substrate concentration while the minor is substrate-concentration independent. In terms of rate of initial luminescence reaction, this increases with the substrate concentration to the power of 1.53 and shows no signs of saturation up to 10 μM coelenterazine. Finally, we found that the inactivated form of the enzyme has a larger apparent size in both size exclusion chromatography and SDS-PAGE analysis and shows a fluorescence peak at 410 nm when excited at 333 nm. These findings indicate that the “flash” kinetics in *Gaussia* luciferase are caused by an irreversible covalent binding to a derivative of the substrate during the reaction.

## Introduction

Bioluminescence is found in a wide variety of organisms ranging from insects and fungi to microorganisms but is most prevalent in marine environments (Martini et Haddock, 2017). *Gaussia princeps* is a deep-sea copepod that lives in the mesopelagic zone and has attracted particular interest for its secreted bioluminescence. In response to various stimuli, it excretes a luminous blue liquid into the surrounding water that is characterized by a rapid increase in intensity followed by a slower decay over 30-80 seconds (Barnes et Case, 1972). It is hypothesized that this allows the copepod to e.g. escape a potential predator. The *Gaussia* luciferase (GLuc) is one of the brightest luminescent proteins found to date (Goerke et al., 2008), despite its size of only 18.2 kDa (after cleavage of its signal peptide). GLuc displays a fairly narrow substrate specificity with coelenterazine (CTZ) being its preferred substrate (Inouye et al., 2013). The enzyme-catalyzed reaction requires oxygen, but no other cofactors. Apart from CO_2_ and light the main CTZ-derived product has not been experimentally determined, but is presumed to be coelenteramide, as in other CTZ utilizing luciferases (Markova et al., 2019). In keeping with its extracellular location, GLuc contains 5 disulfide bridges (Rathnayaka et al., 2011).

GLuc has been developed as a tool in several applications ranging from reporter of gene expression, in bacteria and mammals to split-protein-complementation assays (Maguire et al., 2009; Kim et al., 2011; Remy et Michnick, 2006; Markova et al., 2019). In particular, it has been promoted as a sensitive and useful tool for studying gene expression, systems of secretion (Tannous, 2009) or for monitoring disulfide bond formation (Yu et al., 2018). Due to the reducing conditions in the cytosol of *E. coli* and lack of disulfide isomerases, expression here typically gives very low yields of correctly folded protein. While some of these issues have been addressed by addition of solubility enhancing tags in conjunction with expression at low temperature (Rathnayaka et al., 2011), the best expression systems have been based on secretion from eukaryotic cells (Larionova et al., 2018).

Sequence analysis shows no homology of GLuc to the well-known luciferases from firefly and *Renilla reniformis* and essentially nothing is known about its tertiary structure, except that the primary structure reveals a putative but distinctive two-domain structure. The two domains contain 59 % similar and 35 % identical amino acids over a range of approximately 75 amino acids, in particular with four highly conserved cysteine residues (Wu et al., 2015). This suggest that the conserved cysteines form intra-domain disulfides, that predate a gene duplication event. On the other hand, the fifth disulfide, which is not conserved between the domains, is likely to accommodate an inter-domain disulfide. Several investigations suggest that the two domains have activity individually, albeit significantly reduced, when synthesized as truncated proteins, (Inouye et Sahara, 2008; Hunt et al., 2015). The first domain is preceded by a non-conserved N-terminal region, which is dispensable for activity in the homologous luciferase from *Metridia longa* (Markova et al., 2012).

In line with the early characterization of *G. princeps* luminescence (Barnes et Case, 1972), the activity of GLuc has sometimes been characterized as “flash” kinetics. We set out to try to understand the enzymatic mechanism behind the flash behavior and in the present work we report an investigation of GLuc and its turnover of the substrate CTZ.

We have expressed GLuc in *E. coli* using the so-called CyDisCo system (Gaciarz et al., 2016), in which a yeast mitochondrial thiol oxidase (Erv1p) and the human protein disulfide isomerase (hPDI) are co-expressed with GLuc to help form and shuffle disulfide bonds during folding. Based on preparations of homogeneous and monomeric, correctly folded protein, we here show that the rapid decay of the GLuc light signal (“flash” kinetics) is almost entirely caused by inactivation of GLuc and not substrate depletion, at least within the range of CTZ concentrations commonly used in various applications. We found that this inactivation of the enzyme, presumably by formation of an adduct, is not immediate but takes place after several rounds of catalysis (as little as 10-30 at low substrate concentrations), and that total light output varies with the substrate concentration in a non-linear fashion. We anticipate that these observations are likely to impact the way that GLuc is utilized as a reporter enzyme in the future.

## Results

### Expression in *E. coli* using a helper plasmid encoding protein disulfide isomerase and thiol oxidase

Initially, we were inspired by an expression system previously described (Rathnayaka et al., 2011) that was reported to give good yields and a native-like disulfide bond pattern (as determined by HPLC and CD spectroscopy). We modeled an expression plasmid construct on this system, which most importantly contained an N-terminal His_6_ tag and a C-terminal Solubility Enhancement Peptide (SEP-tag; DDDGDDDGDDDG). We expressed this GLuc reading frame from a T7 promotor, in BL21(DE3) at 37 °C and purified the protein by immobilized metal affinity chromatography (IMAC). To estimate the formation of correct disulfide bonds in the IMAC-purified protein we compared the mobility on SDS-PAGE under reducing vs non-reducing conditions (**Fig 1**). Under reducing conditions much of the protein was found in a defined band consistent with the size of the tagged protein (**Fig 1, lane 3**), however, under non-reducing conditions, we observed that nearly all the purified material was found in non-homogenous high-molecular mass disulfide-linked complexes (HMDC’s) (**Fig 1, lane 5**). Curiously, the HMDC’s showed no sign of aggregation or precipitation and displayed little to no enzymatic activity (**SI fig 5**).

**Figure 1:**
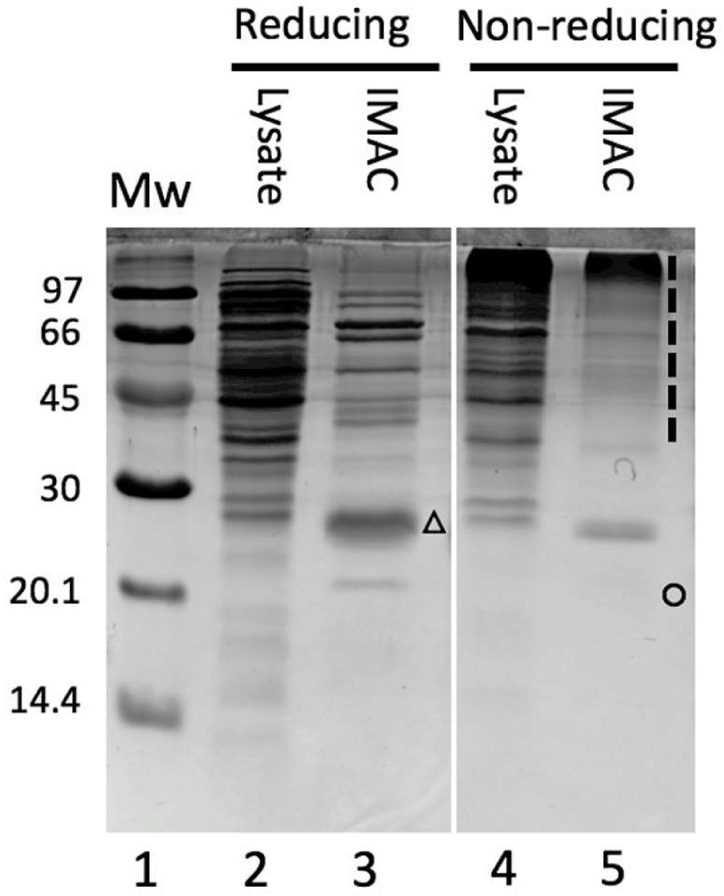
SDS-PAGE of GLuc under reducing and non-reducing conditions, Cell lysate and pooled IMAC eluate corresponding to equal volumes of original culture was run under reducing and non-reducing conditions as indicated. Under non-reducing conditions almost all of the GLuc is found in non-homogenous high-molecular mass disulfide-linked complexes (dashed line, lane 5), which collapse to a single band under reducing conditions (triangle, lane 3). The faster moving band of correctly folded, more compact GLuc under non-reducing conditions is barely visible here (circle), but can be better seen in figure 2. The full gel can be found in the supplementary information (**SI fig 4**).

To reduce the formation of the HDMC’s and increase the yield of monomeric active GLuc we took advantage of the CyDisCo system, in which a plasmid (pMJS205) encoding the yeast mitochondrial thiol oxidase (Erv1p) and human protein disulfide isomerase (hPDI) allows for co-expression of the genes for these enzymes with the gene encoding a disulfide-containing protein of interest (Gaciarz et al., 2016). To study the influence of the CyDisCo system on the proper formation of disulfide bridges in GLuc, we compared protein synthesis in BL21(DE3) cells (**Fig 2a**) with that of this strain previously transformed with pMJS205 (**Fig 2b**).

**Figure 2:**
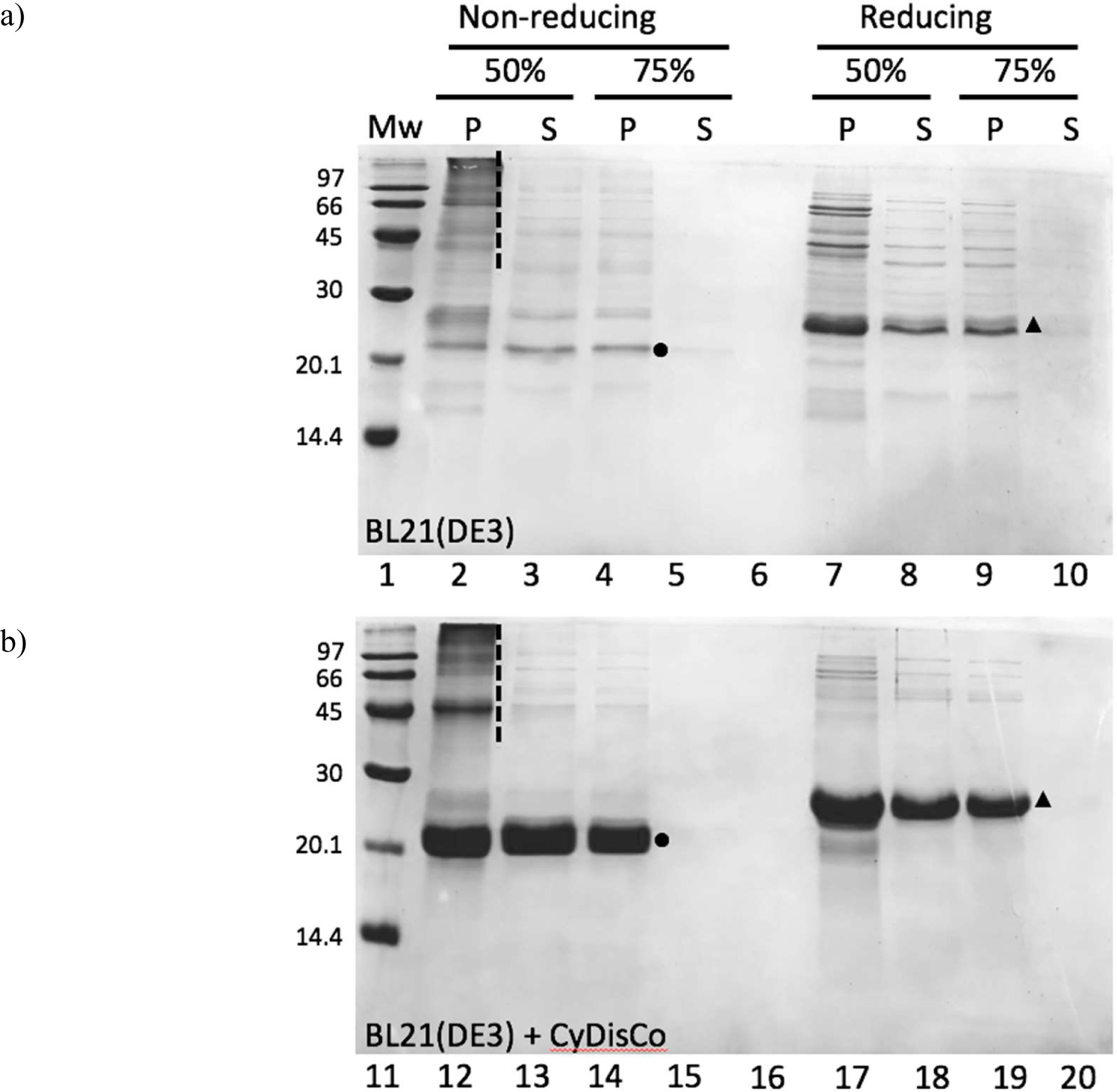
CyDisCo increases the yield of correctly disulfide-linked GLuc and ammonium sulfate precipitation aids the removal of HMDC’s (dashed lines). SDS-PAGE gels showing aliquots from IMAC purified material from equal volumes of culture producing GLuc in the absence (a) and presence (b) of CyDisCo plasmid. Without CyDisCo almost all GLuc is found in HMDC’s (lane 2). While there is still a lot of HMDC in the cells with CyDisCo (lane 12), the fraction of correctly disulfide-linked material is much higher. Correctly folded and disulfide-bonded GLuc (circle) clearly migrates faster than the more extended conformation with reduced disulfide bonds (triangle). The HMDC’s are efficiently removed in the 50 % ammonium sulfate supernatant and precipitated at 75 % ammonium sulfate. Lanes 6 and 16 were left empty to avoid DTT cross-reaction.

A 50 % to 75 % ammonium sulfate fractionation proved highly efficient for separating the HMDC’s and monomeric GLuc. Several interesting observations came from this analysis. In the absence of the CyDisCo plasmid, we found a large amount of heterogeneous HMDC’s under non-reducing conditions (**Fig 2a, lane 2**). That much of this was indeed GLuc is seen from the compression into one band with an apparent molecular mass of 26 kD under reducing conditions (**Fig 2a, lane 7**). Comparing the same AMS fraction on material from the CyDisCo strain, a strong band corresponding to an apparent molecular mass of about 22 kDa appeared on non-reducing SDS-PAGE (**Fig 2b, lane 2**), which shifted upwards to an apparent molecular mass of 26 kDa upon reduction (**Fig 2b, lane 6**), indicating that under non-reducing conditions GLuc has a more compact structure held together by intramolecular disulphide bonds. A similar shift was described earlier for GLuc without a SEP-tag (Goerke et al., 2008). Comparing the 50 % AMS pellet and supernatant, we found that this quite efficiently separated away most of the HMDC’s (most clearly seen in **lanes 2 and 3 of Fig 2b**). Finally, we found that monomeric GLuc was efficiently precipitated at 75 % AMS (**Fig 2, lanes 4 and 14**) leaving essentially no protein in the supernatant (**Fig 2, lanes 5 and 15**).

Although excellent work has been carried out on *in vitro* formation of disulfide bonds in GLuc (Yu et al., 2018) and on cell-free expression of correctly folded monomers (Goerke et al., 2008), the issue of correct disulfide bond formation has not received enough attention in the literature. Non-reducing SDS-PAGE constitutes a very sensitive method for distinguishing both the extended reduced form and HMDC’s from the disulfide linked monomeric form of GLuc.

The term “flash” kinetics and the GLuc reaction

**Figure 3:**
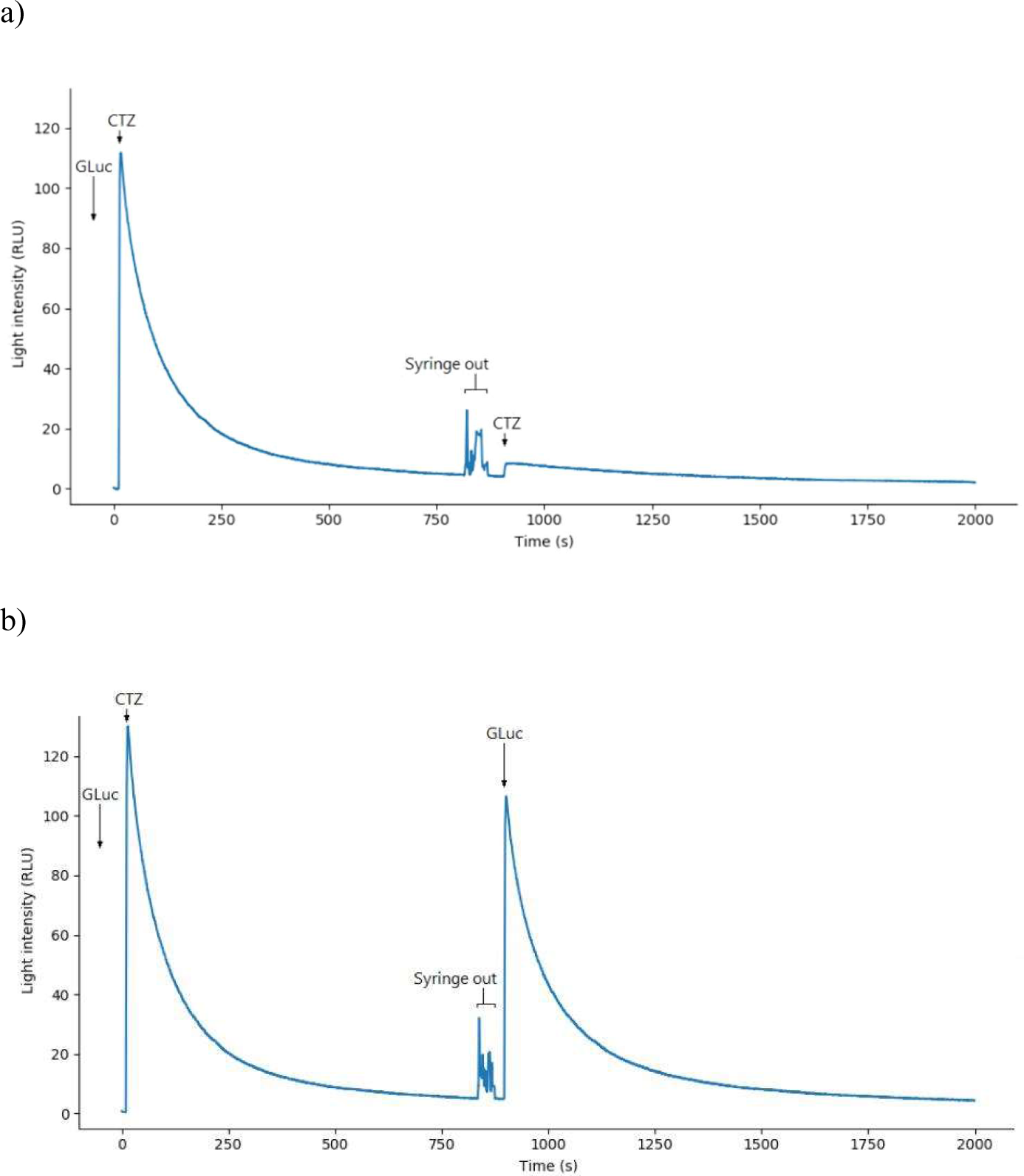
GLuc inactivates during catalysis. **Panel A. Inactivation of GLuc during the reaction**. At t = 10 s, 20 µL CTZ in isopropanol was injected into 2 mL 5.4 nM GLuc to a final concentration of 2.5 μM CTZ, resulting in a peak in light emission. Irregular spikes between 800 s and 900 s were due to the syringe being taken out for the second injection. At 900 s, the same amount of either CTZ (a) or GLuc (b) was added via another 20 µL injection. The lack of light signal when fresh substrate was injected (a) showed that only very little active enzyme was left compared to the beginning of the reaction. The high spike upon injection of fresh enzyme indicated that substrate was still present after the first reaction.

A typical recording of activity in the GLuc assay is shown in **figure 3** where 20 µL of 0.25 mM CTZ is injected into a 2 mL solution containing 5.4 nM GLuc. In this recording an initial steady state rate of photon production is expected to be seen as a constant level of luminescence. However, such a steady state rate was never observed with the substrate concentrations used in the present work and has not been reported previously by others under any conditions. The GLuc kinetics of luminescence are characterized by a very sharp peak that decays over time. Therefore, the best estimate of an initial rate is the height of the luminescence peak immediately after mixing (Fig 3).

In simple enzyme kinetics, an enzyme catalyzes the conversion of a substrate, S, to product, P, by cycling as outlined in **scheme 1** below.

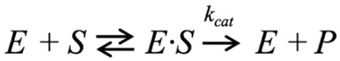

*Scheme 1: Simple enzyme mechanism where the enzyme binds substrate, undergoes reaction and releases the product to form free enzyme*.

A reasonable assumption is that the GLuc reaction with CTZ as substrate is irreversible and only ceases when essentially all substrate has been converted to product. Thus, the most straightforward explanation for the rapidly decreasing activity seen in the first part of the reaction in **figure 3a** is that substrate is completely turned over within 10-15 min. and the addition of more substrate should result in further product formation. However, an additional injection of substrate at t = 900 s, in an amount equal to that injected at t = 10 s, gave virtually no peak (**Fig 3a**), indicating a loss of GLuc activity after the initial substrate addition. Also, if turnover of the entire amount of substrate had not been achieved in the first luminescence peak because of GLuc inactivation, the addition of more GLuc to the incubation should demonstrate this. The latter was exactly what was found (**Fig. 3b**) in an experiment similar to that in **figure 3a** where GLuc injection replaced the addition of a second round of substrate. Based on this, we conclude that GLuc activity ceases before the reaction completes and therefore it is meaningless to apply a general steady state kinetic model to the reaction of GLuc with CTZ. It should be added that incubation of GLuc for 20 min at assay temperature in the absence of substrate resulted in no significant decrease in activity (data not shown).

From the above results it was evident that GLuc was inactivated during turnover of substrate but it was not revealed whether GLuc is inactivated immediately by reacting with the substrate or whether in fact GLuc turns substrate over multiple times before becoming inactivated. For luciferases, the flux of photons from the reaction is equivalent to the rate of the reaction and the integrated photon count over time equivalent to product formation. However, since luminescence output cannot easily be quantified in absolute terms an alternative is to determine the amount of substrate used in the reaction. The amount of remaining CTZ was determined using NanoLuc (England et al., 2016) for which CTZ is also a substrate with a K_m_ of 0.83 μM (**SI fig 7**) and CTZ concentrations could easily be calculated from the NanoLuc light output using a standard curve. An example of determining the remaining CTZ concentration in an incubation after inactivation of GLuc is shown in **figure 4**. CTZ decays spontaneously under the reaction conditions used in the assay with a decay rate determined to be (2.8 ± 0.3) ·10^−4^ s^−1^ (**SI fig 8**), which was included in the calculation of free substrate from experiments performed as in **figure 4**.

**Figure 4:**
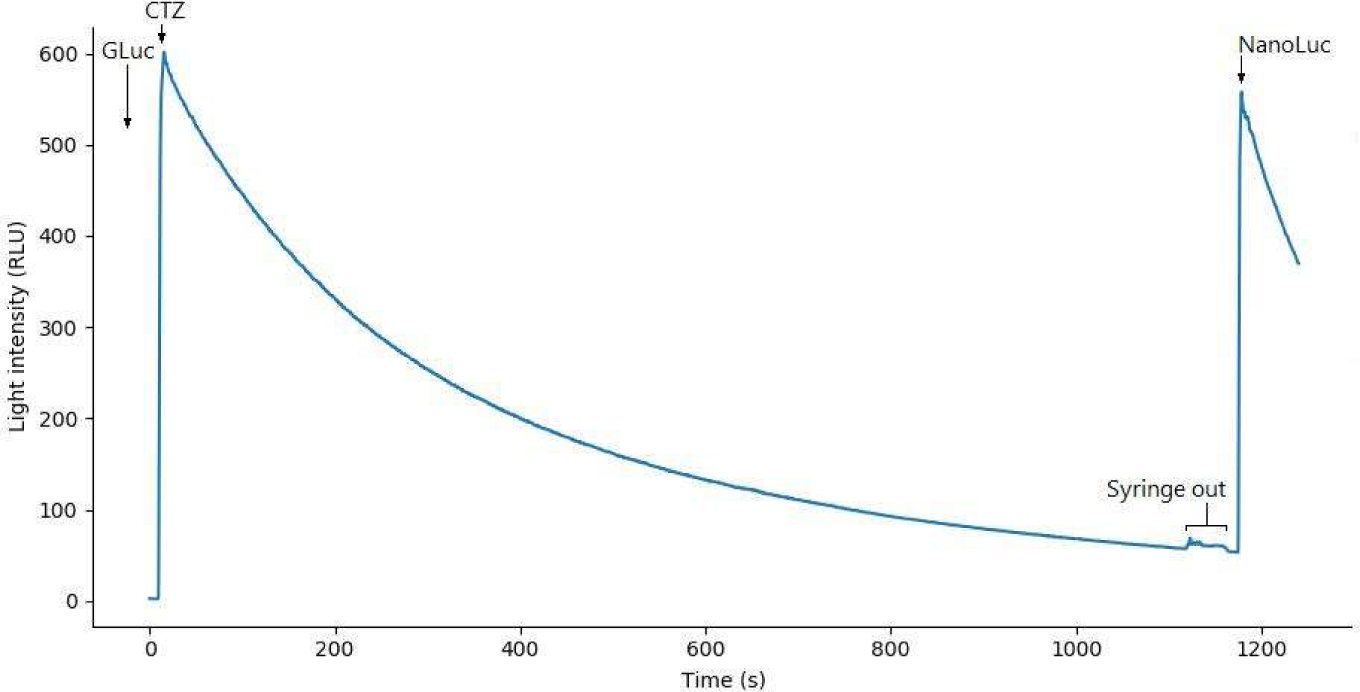
Determination of remaining CTZ after GLuc inactivation. At t = 10 s, 20 µL CTZ in isopropanol was injected into assay buffer containing GLuc, resulting in a total volume of 2 mL with 0.86 µM CTZ and 26.6 nM GLuc and a spike of light emission. Irregular spikes between 1120 s and 1160 s are due to the syringe being taken out. At 1180 s, 20 µl of NanoLuc in assay buffer was injected to a total NanoLuc concentration of 0.2 nM. From the height of the NanoLuc peak we determined how much CTZ was present at the end of the first reaction. Because of the low CTZ concentration after reaction with GLuc, it is rapidly depleted by NanoLuc, resulting in a sharp decline in signal. We checked in a separate experiment that NanoLuc was not inactivated by reacting with CTZ (data not shown).

**Figure 5:**
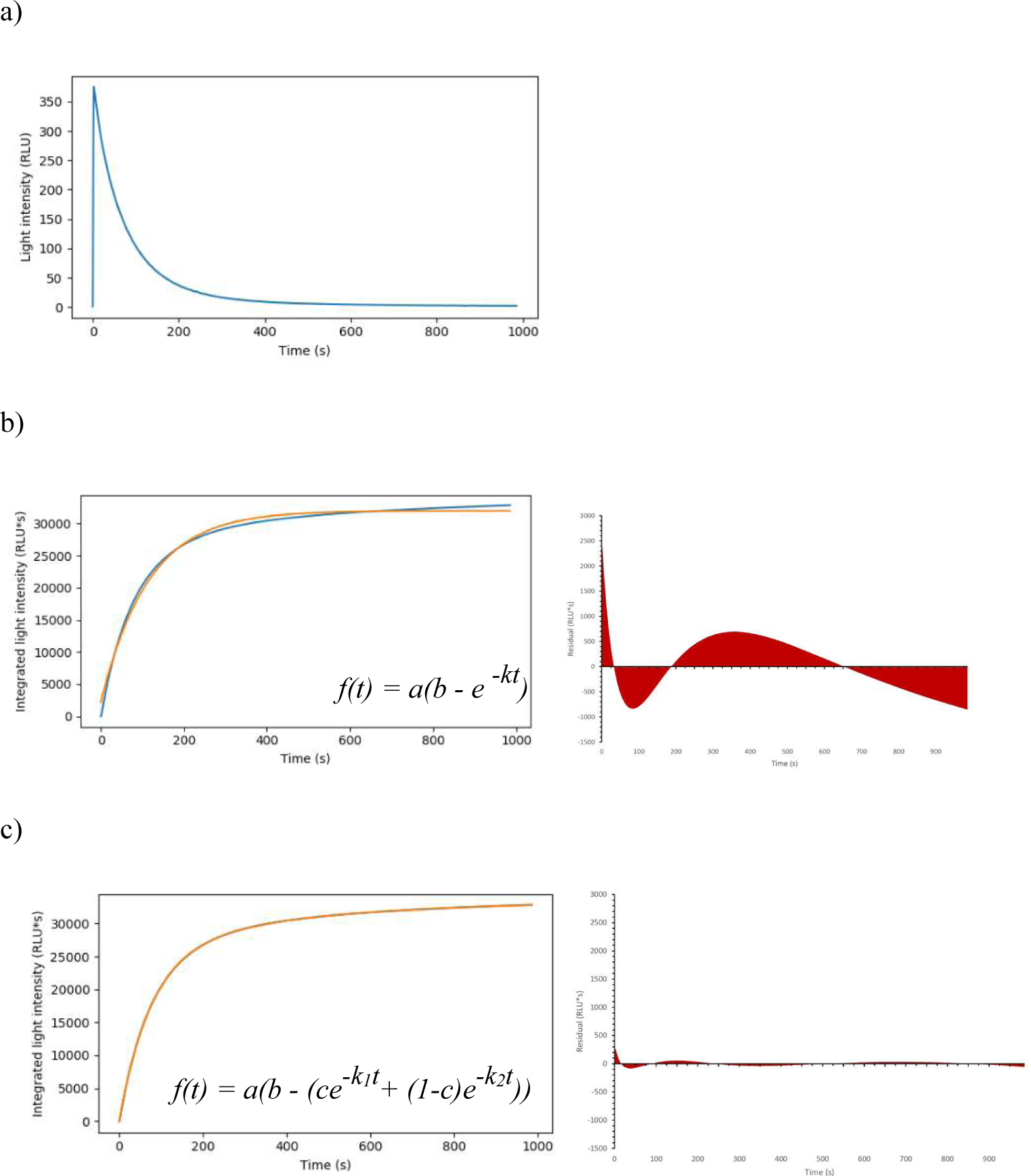
Conversion of luminescence to product formation. A representative assay was made with 0.3 nM Gluc and 3.16 µM CTZ in a reaction volume of 2 ml. a) Light signal after subtraction of background and setting t = 0 at the start of the peak. b) Integrated light signal (blue) onto which a single exponential (orange) was fitted. A considerable residual is left. c) Same data as b (blue) fitted with double exponential (orange). The fit was exceedingly accurate and as seen from almost complete coverage of the data by the fit and the even residual pattern.

**Figure 6:**
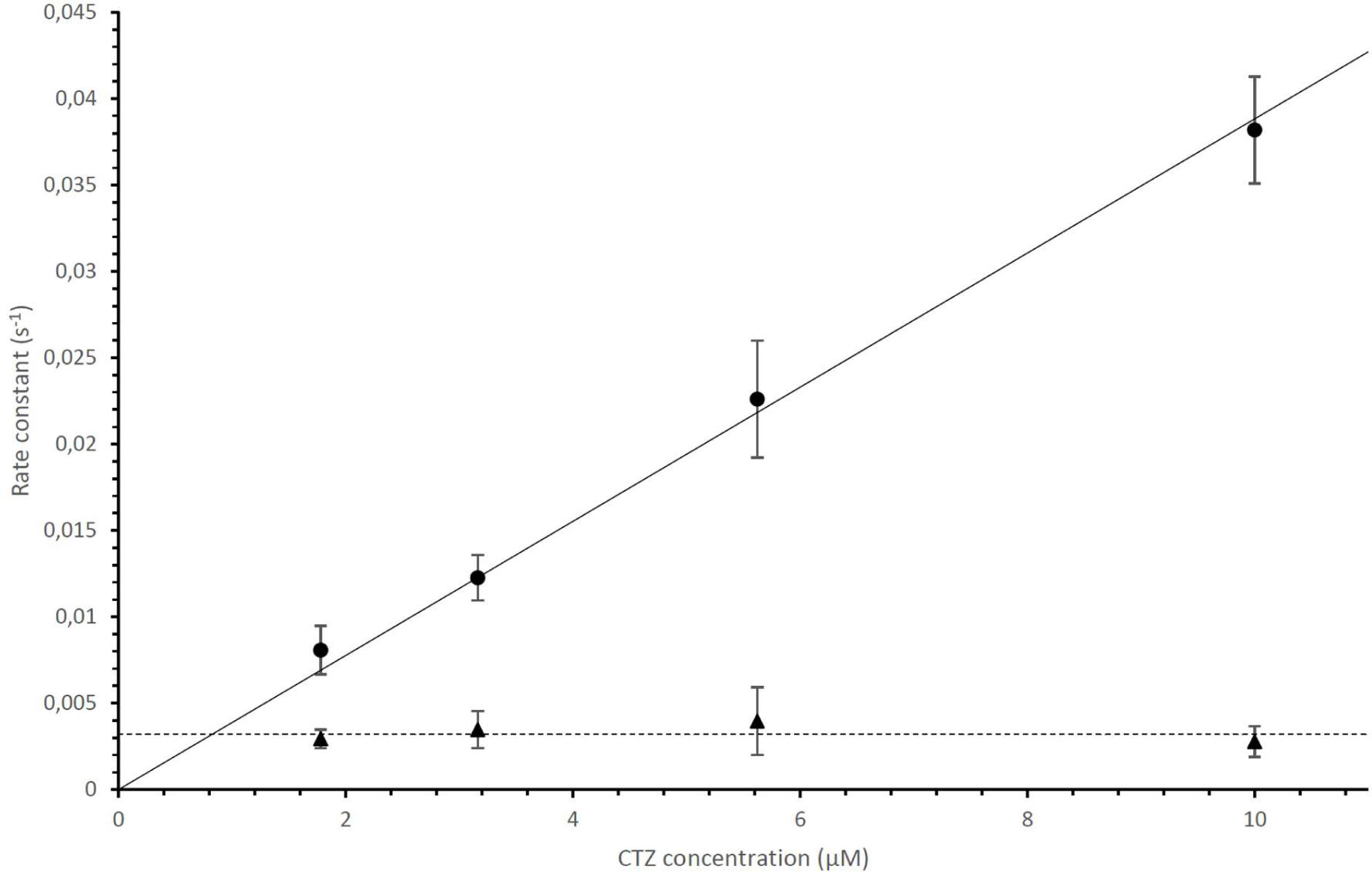
Parameters k1 and k2 of 17 different combinations of CTZ and GLuc concentrations to Eq 2. Data points were an average of 9, 11, 13 and 8 measurements, respectively (low to high CTZ concentration), with GLuc concentrations ranging from 0.051 nM to 0.93 nM; error bars indicating the standard deviation. The slope of the substrate-dependent rate, which we defined as k1, was 3.9·10^−3^ s^−1^µM^−1^ (full line), equivalent to a first order rate constant. The other rate constant, k2, was independent of substrate concentration, with a value of 0.0032 s^−1^ (dashed line). For CTZ concentrations below ∼1.5 μM, the two rate constants k1 and k2 became indistinguishable.

**Figure 7:**
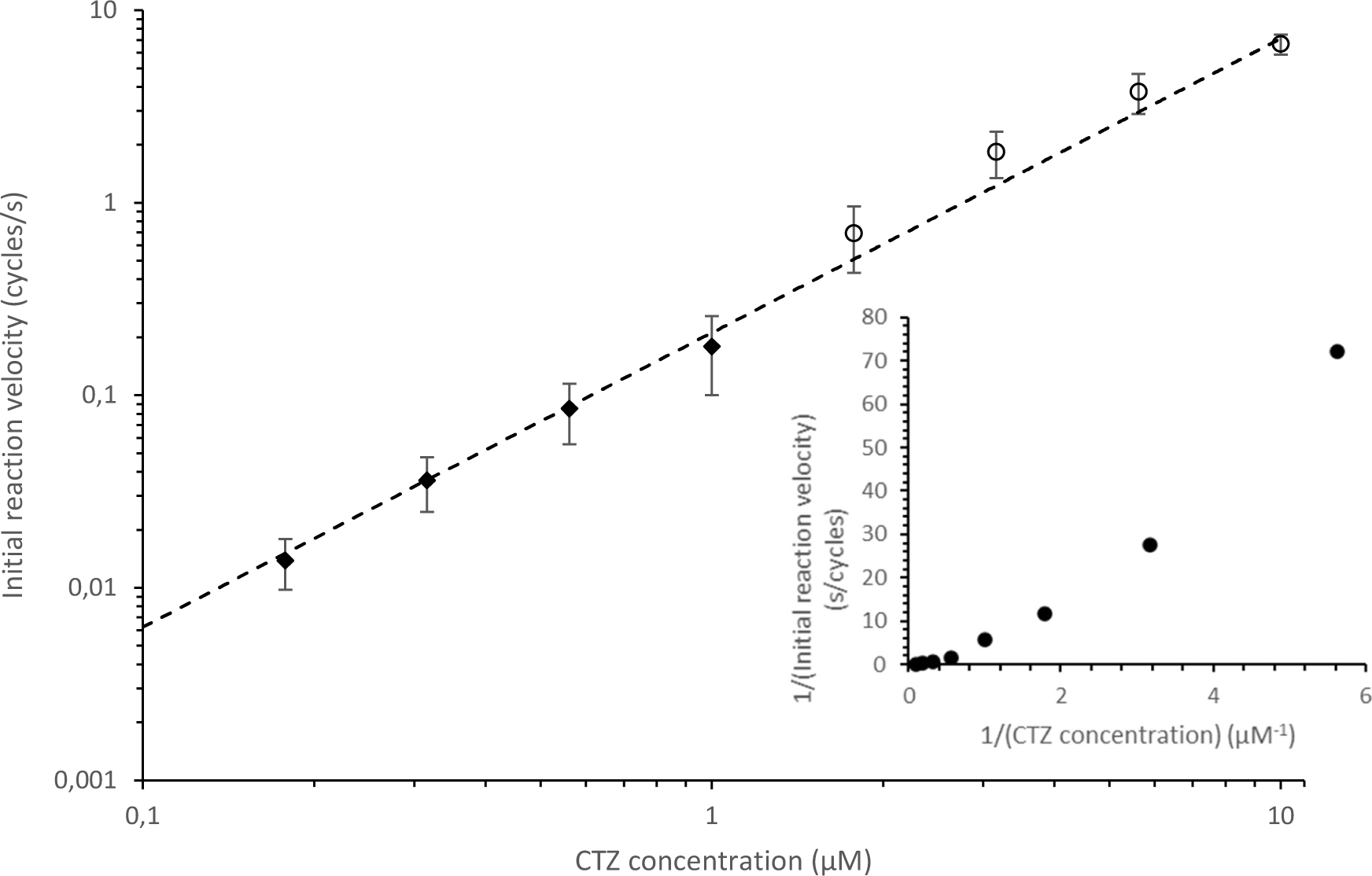
Initial rate versus [CTZ] calculated from fits to equation 1 (black diamonds) or 2 (open circles) and normalized for GLuc concentration. 37 different combinations of CTZ and GLuc concentration were measured. GLuc concentrations ranged from 0.051 nM to 16.2 nM. Each data point is an average of 9-14 measurements. Error bars indicate the standard deviation. The dashed trendline represents 0.15·[S]^1.5^ suggesting that [CTZ] acts co-operatively with respect to catalytic turnover. Between 0.2 and 10 μM CTZ we find no obvious sign of saturation. Insert: both axes reciprocal like a traditional Lineweaver-Burk plot.

**Figure 8:**
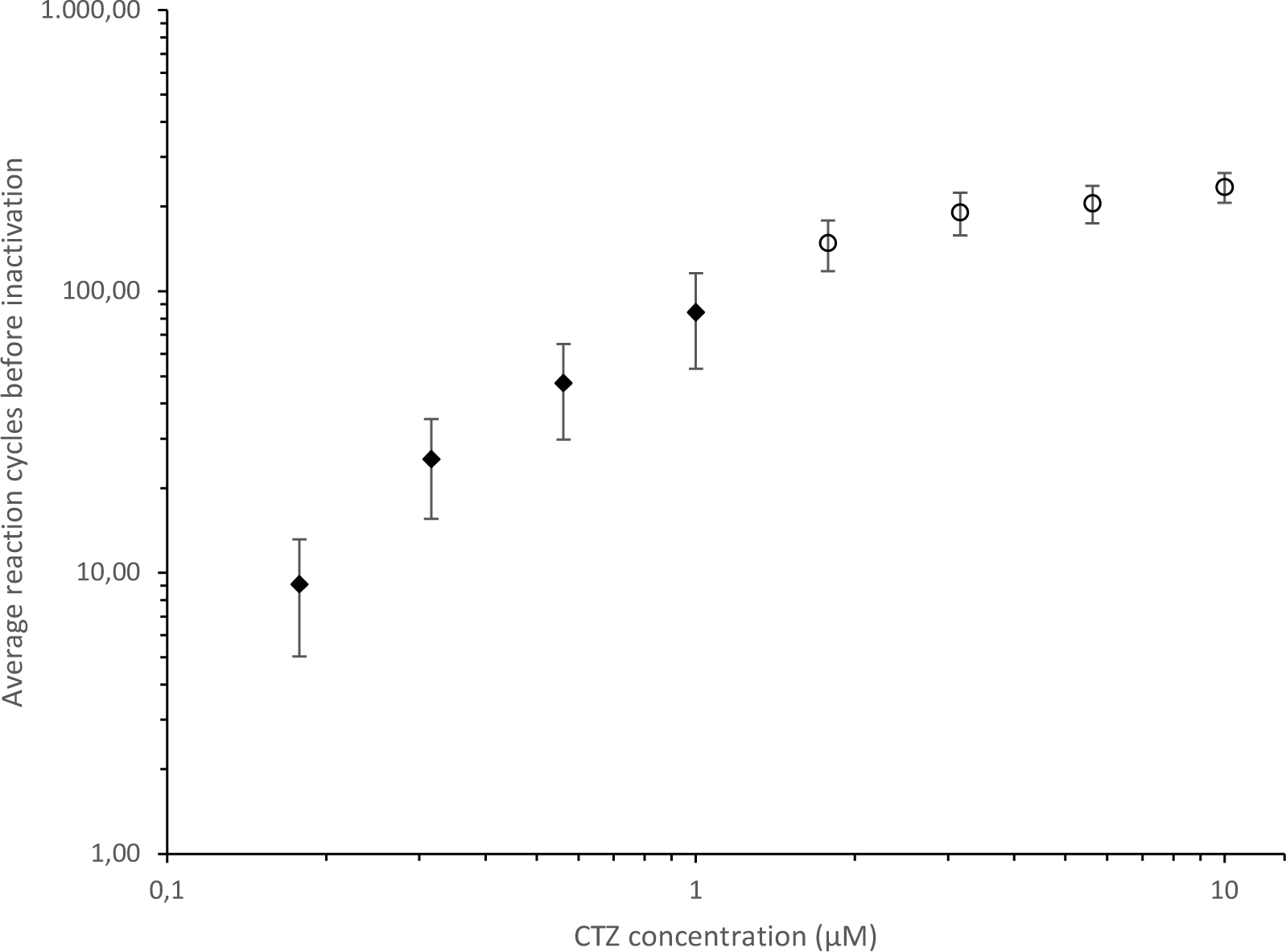
Total number of cycles performed by GLuc is dependent on substrate concentration. Assays for 33 different combinations of CTZ and GLuc concentrations, in which substrate was not used up. The number of cycles was calculated from parameter a from Eq 1 (black diamonds) or Eq 2 (open circles), which is an estimate of the total integral at infinite time, after normalizing it for enzyme concentration. Each datapoint is an average of 7-13 measurements. Error bars indicate the standard deviation. GLuc concentrations ranged from 0.051 nM to 16.2 nM.

From the NanoLuc signal intensity corrected for the spontaneous decay of CTZ, the remaining CTZ from the reaction with GLuc was calculated and from there the number of turnovers performed by the enzyme. For the assay shown in **figure 4**, the number of turnovers was approximately 20. From several such assays, we made a standard curve relating total light output to amount of substrate used. (**SI fig 9**)

**Figure 9:**
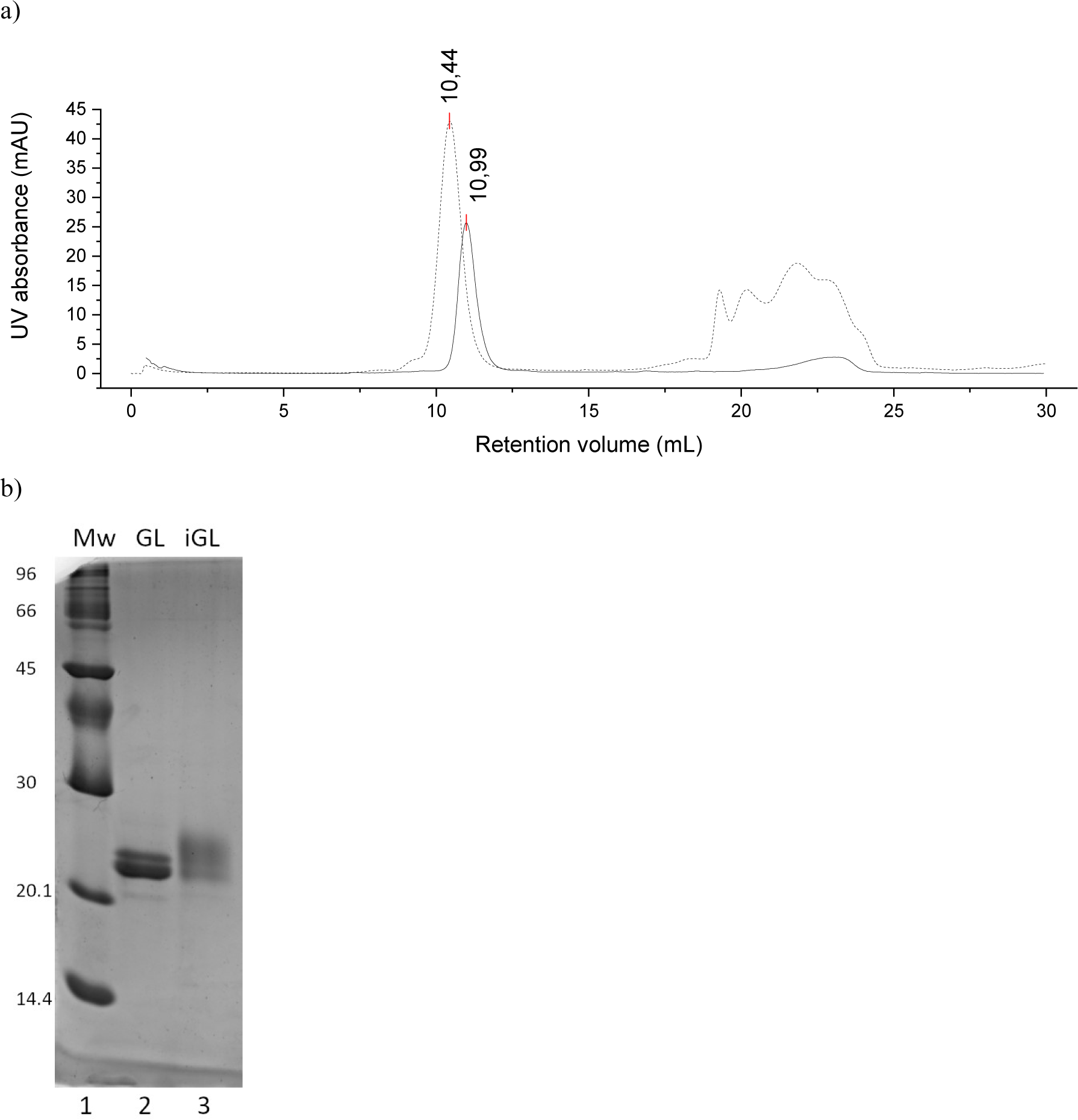
After reaction with CTZ, GLuc had a slightly larger size as measured by size exclusion chromatography (a) and SDS-PAGE (b). a) Active GLuc (solid line) eluted at 10.99 mL on a GE Healthcare Superdex75 10/30 column, whereas after reacting with an excess of CTZ (iGLuc) it eluted at 10.44 mL (dashed line). From a protein standard calibration, this was calculated to equivalent to a shift in apparent molecular weight from 35.3 kDa to 39.5 kDa. b) Equal volumes of the peak fractions of these size exclusions were separated on non-reducing SDS-PAGE. The iGLuc (iGL) shifted upwards relative to unreacted GLuc (GL) and formed a less defined band. The shift persisted under reducing conditions (data not shown).

### Characterization of the inactivation kinetics of GLuc

The shape of the luminescence peak from the GLuc reaction with CTZ appeared to vary with the enzyme and substrate concentrations used. From literature, it is known that the peak height, as a measure of initial velocity *v*_*0*_, increases linearly with enzyme concentration, but that the peak height when varying the CTZ concentration results in a saturation curve displaying positive cooperativity (Larionova et al., 2018; Verhaegen et Christopoulos, 2002). To study GLuc in a conventional enzyme kinetics framework with progress curves of product formation over time, we integrated the light signal to represent product formation (**Fig 5b**).

Datasets were recorded with eight different substrate concentrations ranging from 0.18 µM to 10 µM and 12 different enzyme concentrations ranging from 51 pM to 28.7 nM and the integrated signals were fit to single and double exponential equations. A single exponential (**Eq 1, see methods**) was sufficient for fitting data recorded with CTZ concentrations up to around 2 μM, but for data from assays with above 2 µM CTZ a double exponential equation (**Eq 2, see methods**) was required to describe the integrated luminescence progress curve. The typical improvement when using a double exponential for this data instead of a single, is shown in **figure 5b and c**. In none of the experiments was [O_2_], which typically is at 200-300 μM in buffers at equilibrium with the ambient atmosphere, expected to change significantly during reaction as CTZ never exceeded 10 µM as mentioned above. Larionova et al. (2018) also found signals where a single exponential did not suffice for a good fit at higher substrate concentrations.

The substrate concentration dependence of k_1_ and k_2_ is shown in **figure 6**. Within substrate concentrations between 1.9 µM and 10 µM, *k*_*1*_ from Eq 2 showed a linear correlation, while *k*_*2*_ was independent of substrate concentration. Under conditions where [GLuc]<<[CTZ] the rate constants for inactivation were not dependent on enzyme concentration (**SI fig 11**). Being linearly dependent on the substrate concentration *k*_*1*_ suggested a bimolecular reaction mechanism with a first-order rate constant of 3.9 µM^−1^s^−1^, while *k*_*2*_ was independent substrate concentration (i.e. zero order), with a value of 0.0032 s^−1^.

As substrate concentrations do not drop significantly under the conditions of the assays represented in **figure 6**, it must be stressed that the rate constants do not represent turnover of substrate, but rather reflect the rate at which the enzyme inactivates. Thus, the decay of luminescence almost solely describes the inactivation of the enzyme.

The initial velocity *v*_*0*_, equivalent to the peak height of the luminescence signal, was determined at different concentrations of CTZ, by extrapolating exponential fits to the peak decay to time zero. In **figure 7** the initial velocities are shown as a function of substrate concentration. Although the enzyme clearly is far from saturated at the highest CTZ concentrations, the curve has a clear non-linear character, increasing with the substrate concentration to the power of 1.5 and only just starting to become more linear at 10 µM CTZ. We will come back to this cooperative effect in the discussion section. As described above, substrate depletion was ruled out as the cause of the non-linear shape.

Finally, the average number of cycles of CTZ turnover before GLuc inactivation was calculated for each of the substrate concentrations in the range given above, as described in the method section. A prerequisite for this calculation is that the substrate is not depleted during the reaction, so to ensure that the substrate concentration was reasonably constant, assays in which more than half of the substrate was used up were eliminated.

### GLuc inactivation is due to a covalent modification

To look more into the nature of the substrate-dependent inactivation, we studied the properties of GLuc after it was essentially fully inactivated (having less than 0.03 % residual activity) by reaction with CTZ. It turned out that this substrate-inactivated GLuc (which we term iGLuc) displayed a decreased mobility equivalent to about 1 kD in non-reducing SDS-PAGE relative to GLuc (**Fig 9b**). Since the protein was most likely completely denatured by SDS, the additional mass most likely arose from a covalent modification blocking the substrate-binding site and rendering the enzyme inactive. To further characterize iGLuc, we subjected this to size exclusion chromatography (**Fig 9a**) using a Superdex 75 and here we also observed a difference equivalent to an apparent molecular weight increase from 35.3 kDa to 39.5 kDa.

Because of its extensive pi-electron system, significant absorbance and fluorescence was expected for CTZ and its derivatives and if covalently bound to iGLuc this would likely be visible in the fluorescence spectrum of iGLuc. Indeed, upon excitation at 333 nm we observed a strong emission peak at 410 nm (**Fig 10a**). Interestingly, this spectrum did not match CTZ, which had an emission maximum at 520 nm, but resembled more CTZ that had auto-degraded overnight by reaction with oxygen (**Fig 10b, dashed line**). To gain more insight to the nature of iGLuc, we subjected the sample to MALDI-TOF mass spectrometry. While GLuc had the expected molecular mass, the results from iGLuc were difficult to reproduce and inconclusive, possibly due to an inhomogeneous reaction product.

**Figure 10:**
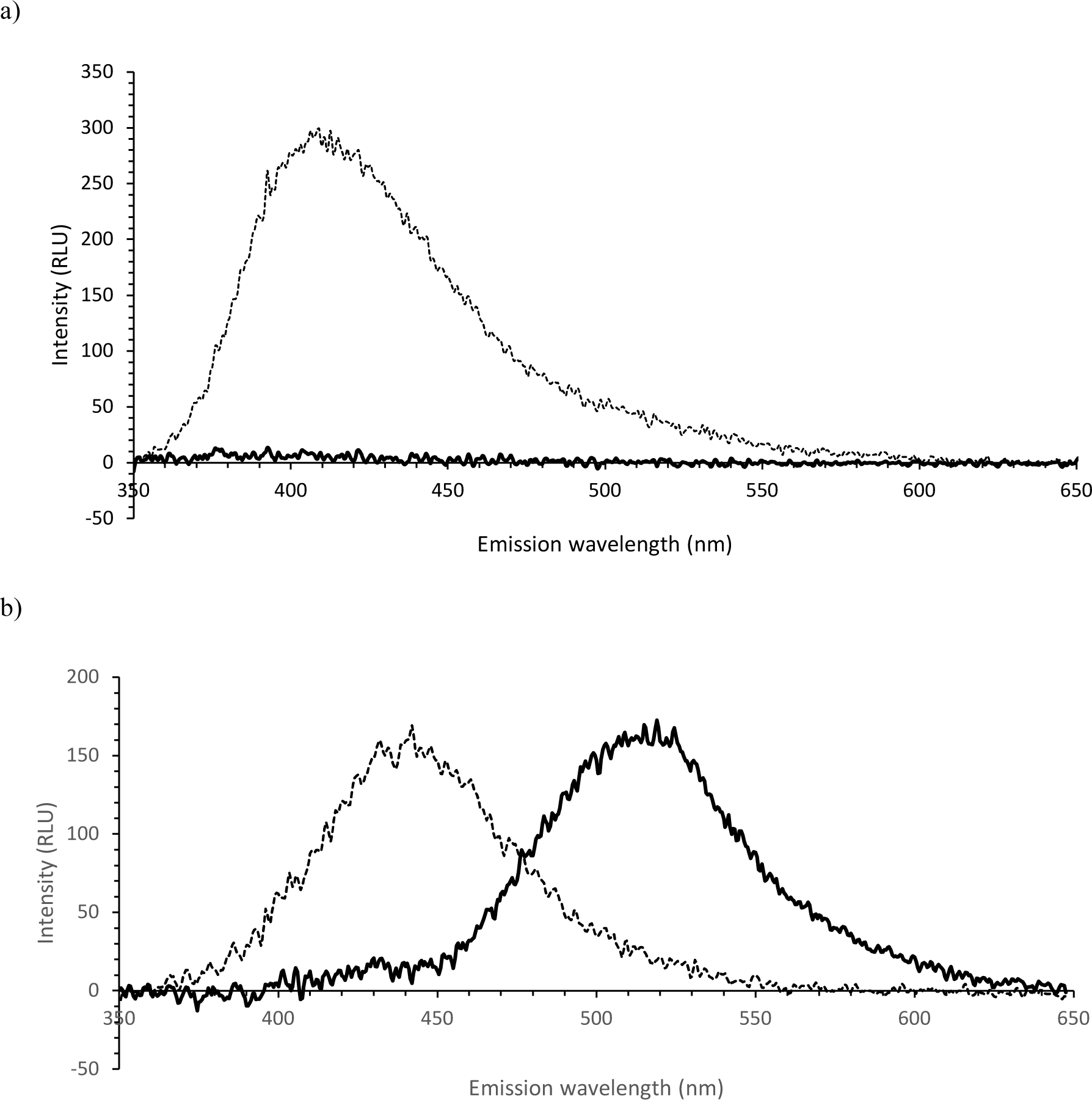
a) Fluorescence spectra excited at 333 nm of iGLuc (dashed line) compared to active GLuc (solid line), both ∼2.2 µM. iGLuc shows a clear emission peak around 410 nm, likely due to a bound CTZ-derivative. b) Fluorescence spectra excited at 333 nm of 6.2 µM fresh CTZ in aqueous buffer at pH 8 (solid line) and the same sample after standing at room temperature overnight (dashed line). A full excitation and emission spectrum of CTZ can be found in the supplementary information (**SI fig 12**).

## Discussion

Despite the wide application of GLuc as a reporter enzyme, the underlying kinetic mechanism behind the term “flash” luciferase has not been investigated in great detail. One reason could be the lack of a good expression system for homogeneous production of this highly disulfide-linked enzyme in *E. coli*, which combined with the surprising observation that the HMDC’s (**Fig 1 and 2**) did not form precipitates should warrant caution. The use of non-reducing SDS-PAGE (**Fig 1 and 2**) was a very sensitive tool not only to distinguish HMDC’s from the disulfide-linked monomer, but also to demonstrate that a homogeneous, monomeric form of GLuc was almost absent in the BL21(DE3) expression system, in our hands. A similar approach using native PAGE has been applied before to GLuc’s homolog in *Metridia longa*, showing that this protein also forms HMDC’s (Borisova, 2008). Using the so-called CyDisCo system, we have shown that introduction of two redox enzymes dramatically increased the yield of monomeric disulfide-linked GLuc, as they allowed more efficient formation and shuffling of disulfides in the cytosol of *E. coli*. The disulfide-linked monomer showed a distinctly higher mobility on non-reducing SDS-PAGE than the corresponding reduced monomer. This compact form of the enzyme eludes as a single active peak on a Superdex75 Size Exclusion Column (**Fig 9a, solid line** and **SI fig 5**), indicating that this constitutes a homogeneous preparation of monomeric protein. The hydrodynamic volume of GLuc is significantly larger than expected for a compact protein, but not large enough to be a dimer. We attribute this large volume in part to the SEP tag, but it may also indicate a generally more dynamic structure of the catalytic domain of this protein. Yields in excess of 5 mg of homogeneous GLuc were routinely obtained per liter culture.

Our experiments showed that the reason that GLuc demonstrates a rapid drop in light emission, even with an excess of substrate and less than 10 % substrate depletion, is that the enzyme is inactivated during the reaction. The inactivation is accompanied by a chemical modification which is most likely a CTZ-derived adduct (**Fig 9 and 10**). To describe the kinetics of the inactivation of GLuc, we found that two exponential rate constants were required for an accurate fit. Both were independent of GLuc concentration and while one was also independent of CTZ concentration, the other showed a first order dependence on CTZ.

The initial velocity increased linearly with the GLuc concentration, but to the power of 1.5 with the CTZ concentration (**Fig 7**). This is unlikely to be an artefact, since especially the initial velocities at lower substrate concentrations would have to be underestimated by up to a factor 10 to match the linear relationship. The value of 1.5 is consistent with a Hill-coefficient of 1.8±0.2 reported previously (Larionova et al., 2018). It has been suggested that this cooperative effect is the result of multiple binding sites (Tzertzinis et al., 2012) or a kinetic cooperativity in which a substrate induced conformational change happens at the same time scale as catalytic turnover (Larionova et al., 2018). Considering the apparent dynamics suggested by the size exclusion chromatography, we favor a kinetic cooperative effect over multiple binding sites in this rather small protein.

It is surprising that the cooperative dependence on substrate concentration does not show up in the decay rates and that the number of reaction cycles before inactivation increases with substrate concentration (**Fig 8**). In a conventional suicide substrate reaction, the normal reaction and inactivation reaction share a reaction intermediate, which results in a fixed inactivation probability in each cycle and thus a fixed average number of reaction cycles (Waley, 1980). If such a reaction would additionally involve cooperativity, this would show up in the reaction velocity as well as the decay rate. This is however not observed, suggesting that there is no common intermediate and the light-emitting reaction and the covalent modification instead follow different pathways, for example as a result of the substrate binding in different orientations.

Photoproteins offer an interesting example of luminescent proteins that covalently bind CTZ after reaction. Obelin is such a photoprotein, a complex of 2-hydroperoxycoelenterazine and apoprotein, which upon addition of Ca^2+^ catalyse a single luminescent reaction. Coelenteramide thus remains tightly associated with Obelin, a complex termed *discharged photoprotein*. Without addition of Ca^2+^, the reaction can also be triggered by heat, resulting a in thermo-inactivated discharged photoprotein with a distinctly different fluorescence spectrum. The fluorescent spectrum of inactivated GLuc shows some similarity to that of thermo-inactivated discharged Obelin (Belogurova et Kudryasheva, 2010), with peaks around the same wavelengths, but with a much larger component at 410 nm compared to 510 nm. This similarity suggests that a similar CTZ-derivative may be accommodated in both iGLuc and thermo-inactivated Obelin. iGLuc shows more intense fluorescence than CTZ or its degradation product in aqueous solution (**Fig 10a and b**), likely due to stabilization of the adduct in iGLuc. The fluorescent properties of iGLuc could open the possibility for its use as a fluorescent reporter, in a similar manner as photoproteins.

We have not been able to find examples of the particular kind of kinetics/inactivation described here in the literature, and one may well wonder how auto-inactivation has survived the natural selection of evolution. We would like to suggest that since the copepod secretes both GLuc and CTZ into the surrounding water, there is only a very short time window for the reaction to occur before dilution will prevent the GLuc:CTZ complex formation. Thus, inactivation of GLuc after a limited number of reaction cycles is not detrimental in the natural context. Evolutionary optimization for the most efficient enzyme – one with a high number of reaction cycles in the first moments of mixing with CTZ – could have led to an enzyme where persistence is of minor importance relative to a high initial light output.

## Concluding remarks

Although a full mechanistic explanation was not reached in this investigation, we can firmly conclude that the rapid decrease in activity seen when assaying this enzyme is an effect of an irreversible inactivation of the enzyme after several reaction cycles. This property is highly unusual and possibly unique and considering the widespread use of this enzyme as a reporter and screening tool, we find it highly relevant for the labs where it is in use. A similar inactivation is likely to take place in GLuc homologs in other copepod species that display flash kinetics.

Finally, we show that the inactivation of GLuc is a likely result of the formation of a covalent link with a CTZ derivative, but further research is necessary to determine the chemical nature of this modification and possibly mitigate it to prolong the lifetime of the enzyme.

## Materials and methods

### Genetic materials

For expression we designed an expression system in a pET-21c vector modeled on an earlier described expression system for GLuc in which the N-terminal signal sequence was replaced by a six-residue histidine-tag and which included a C-terminal Solubility Enhancing Peptide (SEP) tag (Ratnayaka et al., 2011). We modified this system by codon optimizing the open reading frame for *E. coli* and replacing factor X_a_-cleavage sites flanking the central GLuc sequence with TEV-protease cleavage sites (see SI for full sequence information). We are grateful to Yukata Kuroda of Tokyo University of Agriculture and Technology for providing the sequence information for the previously described plasmid (Ratnayaka et al., 2011). The remodeled plasmid (pAKD01) was custom synthesized by Genscript. For proper disulfide bridge formation, we used the CyDisCo system (Gaciarz et al., 2016). This system allows disulfide bond formation in the *E. coli* cytosol as it expresses a polycistronic mRNA encoding the sulfhydryl oxidase Erv1p, and the human Protein Disulfide-Isomerase (hPDI). The plasmid, pMJS205, which also contained the cat-gene for resistance towards chloramphenicol was kindly provided by Lloyd Ruddock, Oulu University, Finland.

### Expression and purification

We tested production of codon optimized GLuc from pAKD01 with His- and Sep-tags in the BL21(DE3) *E. coli* strain both without and with pMJS205 (CyDisCo). For expression, cells were grown in liquid AB-LB medium (Lauritsen et al., 2011), at 37 °C and induced with 0.2 mM IPTG, after which protein was expressed overnight at 37 °C. To prepare the cell-extract for purification, cells were spun down, sonicated in lysis buffer and any leftover DNA was removed by streptomycin precipitation. The protein was purified by immobilized metal affinity chromatography (IMAC). Peak elution fractions were combined and ammonium sulfate at 50 % saturation to precipitate and remove primarily high molecular weight cross-linked GLuc. The supernatant was then adjusted to 75 % ammonium sulfate saturation to precipitate monomeric GLuc. The pellet was finally solubilized in storage buffer and dialyzed against 50 mM Tris buffer pH 8.0 to remove ammonium sulfate. See supplementary information for full details.

### Luminescence assays

All luminescence assays were performed in 2 ml 50 mM Tris buffer, pH 8, with 0.1 gL^−1^ BSA to prevent adhesion of luciferase to the cuvette walls at the very low concentrations frequently used. Light emission at 480 nm was measured using a Perkin Elmer LS55 Luminescence spectrometer with 3 ml stirred cuvette at 25 °C. Unless stated otherwise, background luminescence was recorded and assays were initiated by addition of CTZ to assay buffer containing luciferase. We used a custom designed syringe holder to inject into the cuvette while the lid was closed (**SI fig 3**). To ensure consistent conditions, all CTZ solutions were kept in isopropanol at -20 °C until use. During the preparation of the assay GLuc and CTZ were kept on ice. As higher concentrations of isopropanol significantly inhibit GLuc (**SI fig 6**), the final isopropanol concentration in all assays was 1 %, except when multiple injections were done. During data analysis, the background was subtracted and the time was set to zero at the onset of the reaction. Mixing time was estimated to be up to 10 s.

### Measurement of number of reaction cycles

Because NanoLuc (England et al., 2016) uses CTZ as substrate just like GLuc, but obeys conventional Michaelis-Menten kinetics we used this enzyme to determine residual CTZ concentrations based on a standard curve. The NanoLuc used was prepared as described in the supplementary information. Assays were performed as described above, but light output was measured at 460 nm instead of 480 nm, to optimize for NanoLuc’s emission maximum. In a separate experiment we determined that the GLuc light signal at this wavelength was 1.25-fold lower than at 460 nm. When the GLuc light signal had reached 10 % of the original peak height, 0.2 nM NanoLuc was injected. The CTZ concentration was estimated by comparing the average light signal 10-20 s after injection to a standard curve determined in the same way. The change in CTZ concentration since the beginning of the GLuc reaction was adjusted for the CTZ half-life in assay buffer which was determined to be 41 min. A standard curve was made, relating light output in RLU·s to the amount of CTZ turned over (**SI fig 9**).

### Light signal curve fitting

A dataset of 37 assays was recorded, consisting of combinations of 8 different substrate concentrations ranging from 0.18 µM to 10 µM and 12 different enzyme concentrations ranging from 51 pM to 28.7 nM. To be able to measure over a wide range of CTZ concentrations, lower enzyme concentrations were combined with higher CTZ concentrations. Each assay was performed in triplicate. Signals were then fit to either a single (**Eq 1**) or double (**Eq 2**) exponential equation.

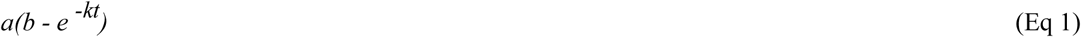

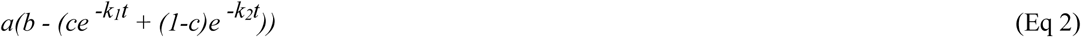

In both the single (**Eq 1** and double (**Eq 2**) exponential equation, parameter *a* determines the asymptote that is approached at infinite time and indicates the maximum light production under the given conditions. *k, k*_*1*_ and *k*_*2*_ are rate constants and *c* is the ratio of the contribution of the two exponentials in the double exponential equation (**Eq 2**). We also added parameter *b*, which corrected the small offset at the beginning of the reaction, when the signal is not yet stable due to mixing and this was usually very close to 1 (**SI fig 10**). For CTZ concentrations above 1.5 µM, we used the double-exponential fit, but for the lower CTZ concentrations we used a single exponential fit, since the rate constants were too close together to be distinguished by the fitting algorithm (**Fig 6**).

### Kinetic analysis

Initial velocities were calculated from a back-extrapolation of an exponential fit of each light signal. Relative light units were converted to reaction velocity in cycles per second using the standard curve made in the NanoLuc experiment described above. The number of reaction cycles before inactivation was calculated from the (extrapolated) integral at infinite time in similar fashion.

### Properties of inactivated GLuc

Inactivated GLuc was prepared by adding 100 µL CTZ in isopropanol to 400 µL assay buffer containing GLuc, to a total CTZ concentration of 0.26 mM CTZ and 13.7 µM GLuc. The mixture was left at room temperature to react overnight and measured less than 0.03 % residual activity the next day, compared to a reference incubated under the same conditions, but with isopropanol only.

Both samples were subjected to size exclusion chromatography on a Superdex75 column. Fractions of 0.5 mL were collected and the two peak fractions of each sample were combined for spectral analysis. Fluorescence spectra were recorded in a 1 mL quartz cuvette on a Perkin Elmer LS55 Luminescence spectrometer. The concentration of active GLuc was 2.2 µM and the concentration of iGLuc similar. Spectra of CTZ were recorded in the same buffer, at a CTZ concentration of 6.2 µM. One spectrum was recorded immediately upon adding CTZ to the buffer, and a second one after the mixture had stood at room temperature overnight and had thus reacted with oxygen in the buffer.

After recording the spectra, 500 µL of GLuc and iGLuc, respectively, was concentrated by spinning at 14000 g for 30 minutes on a preequilibrated microcon10 centrifugal filter. 10 µL of each concentrated sample was analyzed by SDS-PAGE. Another 10 µL was used to generate a mass spectrum on a MALDI-TOF MS system, but the results were inconclusive.

## Acknowledgements

We would like to thank Dr Lloyd Ruddock for kindly providing the CyDisCo plasmid and for helpful communication and to thank Dr. Yutaka Kuroda for providing us with the sequence and plasmid for GLuc that our own construct was based on. Thanks to Benjamin Bjerre for providing us with purified NanoLuc and to Charlotte O’Shea for help with experiments and discussion of results.

## References

Barnes, A. T., & Case, J. F. (1972). Bioluminescence in the mesopelagic copepod, Gaussia princeps (T. Scott). Journal of Experimental Marine Biology and Ecology, 8(1), 53–71.

Belogurova, N. V., & Kudryasheva, N. S. (2010). Discharged photoprotein obelin: fluorescence peculiarities. Journal of photochemistry and photobiology. B, Biology, 101(1), 103–108. https://doi.org/10.1016/j.jphotobiol.2010.07.001

Borisova, V. V., Frank, L. A., Markova, S. V., Burakova, L. P., & Vysotski, E. S. (2008). Recombinant Metridia luciferase isoforms: expression, refolding and applicability for in vitro assay. Photochemical & Photobiological Sciences, 7(9), 1025–1031.

England, C. G., Ehlerding, E. B., & Cai, W. (2016). NanoLuc: a small luciferase is brightening up the field of bioluminescence. Bioconjugate chemistry, 27(5), 1175–1187.

Gaciarz, A., Veijola, J., Uchida, Y., Saaranen, M. J., Wang, C., Hörkkö, S., & Ruddock, L. W. (2016). Systematic screening of soluble expression of antibody fragments in the cytoplasm of E. coli. Microbial cell factories, 15(1), 22.

Goerke, A. R., Loening, A. M., Gambhir, S. S., & Swartz, J. R. (2008). Cell-free metabolic engineering promotes high-level production of bioactive Gaussia princeps luciferase. Metabolic engineering, 10(3-4), s187–200.

Hunt, E. A., Moutsiopoulou, A., Ioannou, S., Ahern, K., Woodward, K., Dikici, E., Daunert, S., & Deo, S. K. (2016). Truncated Variants of Gaussia Luciferase with Tyrosine Linker for Site-Specific Bioconjugate Applications. Scientific reports, 6, 26814. https://doi.org/10.1038/srep26814

Inouye, S., Sahara-Miura, Y., Sato, J. I., Iimori, R., Yoshida, S., & Hosoya, T. (2013). Expression, purification and luminescence properties of coelenterazine-utilizing luciferases from Renilla, Oplophorus and Gaussia: comparison of substrate specificity for C2-modified coelenterazines. Protein Expression and Purification, 88(1), 150–156.

Inouye, S., & Sahara, Y. (2008). Identification of two catalytic domains in a luciferase secreted by the copepod Gaussia princeps. Biochemical and biophysical research communications, 365(1), 96–101. https://doi.org/10.1016/j.bbrc.2007.10.152

Kim, S. B., Suzuki, H., Sato, M., & Tao, H. (2011). Superluminescent variants of marine luciferases for bioassays. Analytical chemistry, 83(22), 8732–8740. https://doi.org/10.1021/ac2021882

Larionova, M. D., Markova, S. V., & Vysotski, E. S. (2018). Bioluminescent and structural features of native folded Gaussia luciferase. Journal of Photochemistry and Photobiology B: Biology, 183, 309–317.

Lauritsen, I., Willemoës, M., Jensen, K. F., Johansson, E., & Harris, P. (2011). Structure of the dimeric form of CTP synthase from Sulfolobus solfataricus. Acta crystallographica. Section F, Structural biology and crystallization communications, 67(Pt 2), 201–208. https://doi.org/10.1107/S1744309110052334

Maguire, C. A., Deliolanis, N. C., Pike, L., Niers, J. M., Tjon-Kon-Fat, L. A., Sena-Esteves, M., & Tannous, B. A. (2009). Gaussia luciferase variant for high-throughput functional screening applications. Analytical chemistry, 81(16), 7102–7106. https://doi.org/10.1021/ac901234r

Markova, S. V., Burakova, L. P., & Vysotski, E. S. (2012). High-active truncated luciferase of copepod Metridia longa. Biochemical and biophysical research communications, 417(1), 98–103. https://doi.org/10.1016/j.bbrc.2011.11.063

Markova, S. V., Larionova, M. D., & Vysotski, E. S. (2019). Shining Light on the Secreted Luciferases of Marine Copepods: Current Knowledge and Applications. Photochemistry and photobiology, 95(3), 705–721. https://doi.org/10.1111/php.13077

Martini, S., & Haddock, S. H. (2017). Quantification of bioluminescence from the surface to the deep sea demonstrates its predominance as an ecological trait. Scientific reports, 7(1), 1–11.

Rathnayaka, T., Tawa, M., Nakamura, T., Sohya, S., Kuwajima, K., Yohda, M., & Kuroda, Y. (2011). Solubilization and folding of a fully active recombinant Gaussia luciferase with native disulfide bonds by using a SEP-Tag. Biochimica et biophysica acta, 1814(12), 1775–1778. https://doi.org/10.1016/j.bbapap.2011.09.001

Remy, I., & Michnick, S. W. (2006). A highly sensitive protein-protein interaction assay based on Gaussia luciferase. Nature methods, 3(12), 977–979. https://doi.org/10.1038/nmeth979

Tannous B. A. (2009). Gaussia luciferase reporter assay for monitoring biological processes in culture and in vivo. Nature protocols, 4(4), 582–591. https://doi.org/10.1038/nprot.2009.28

Tzertzinis, G., Schildkraut, E., & Schildkraut, I. (2012). Substrate cooperativity in marine luciferases. PloS one, 7(6), e40099.

Verhaegen, M., & Christopoulos, T. K. (2002). Recombinant Gaussia luciferase. Overexpression, purification, and analytical application of a bioluminescent reporter for DNA hybridization. Analytical chemistry, 74(17), 4378–4385.

Waley S. G. (1980). Kinetics of suicide substrates. The Biochemical journal, 185(3), 771–773. https://doi.org/10.1042/bj1850771

Wu, N., Rathnayaka, T., & Kuroda, Y. (2015). Bacterial expression and re-engineering of Gaussia princeps luciferase and its use as a reporter protein. Biochimica et Biophysica Acta (BBA)-Proteins and Proteomics, 1854(10), 1392–1399.

Yu, T., Laird, J. R., Prescher, J. A., & Thorpe, C. (2018). Gaussia princeps luciferase: a bioluminescent substrate for oxidative protein folding. Protein science: a publication of the Protein Society, 27(8), 1509–1517. https://doi.org/10.1002/pro.3433

