## Supplementary information for "Flash properties of Gaussia Luciferase are the result of covalent inhibition after a limited number of cycles"

### Supplementary materials and methods

#### Buffers and media

**AB-LB medium**

| | |
|---|---|
| Bacto-peptone | 10 g $l^{-1}$ |
| Bacto-yeast extract | 5 g $l^{-1}$ |
| NaCl | 5 g $l^{-1}$ |
| $MgCl_2$ | 2 mM |
| $CaCl_2$ | 0.1 mM |
| $FeCl_3$ | 3 µM |
| After autoclaving: | |
| $(NH_4)SO_4$ | 2 g $l^{-1}$ |
| $Na_2HPO_4 \cdot 12\ H_2O$ | 15 g $l^{-1}$ |
| $KH_2PO_4$ | 3 g $l^{-1}$ |
| NaCl | 3 g $l^{-1}$ |
| Glucose | 0.2 % |

**Concentrations of antibiotics** were the following unless mentioned otherwise:

| | |
|---|---|
| Chloramphenicol | 50 µg $ml^{-1}$ |
| Ampicillin | 100 µg $l^{-1}$ |

**Lysis buffer**

| | |
|---|---|
| Tris | 50 mM |
| NaCl | 150 mM |
| Imidazole | 10 mM |
| Triton X-100 | 1% |

Adjusted to pH 8.0 with HCl

Filtered through a 0.22 µm vacuum filter

**IMAC wash buffer**

| | |
|---|---|
| Tris | 50 mM |
| NaCl | 150 mM |
| Imidazole | 20 mM |

Adjusted to pH 8.0 with HCl

Filtered through a 0.22 µm vacuum filter

**IMAC elution buffer**

| | |
|---|---|
| Tris | 50 mM |
| NaCl | 150 mM |
| Imidazole | 600 mM |

Adjusted to pH 8.0 with HCl

Filtered through a 0.22 µm vacuum filter

**Assay buffer/storage buffer**

| | |
|---|---|
| Tris | 50 mM |
| BSA | 0.1 g $l^{-1}$ |

Adjusted to pH 8.0 with HCl

Filtered through a 0.22 µm vacuum filter

During assays there was an additional isopropanol concentration of 1 %, because CTZ was stored and diluted in this solvent.

#### Genetic materials

In our experiments, we used GLuc with the same sequence as the Kuroda group. This sequence differs from the canonical wild type by two amino acids (E100A and G103R). As seen in **supplementary figure 1**, we used two tags: a HIS-tag for nickel affinity purification at the N-terminus and a solubility enhancement (SEP) tag at the C-terminus. Both tags were separated from the rest of the sequence by a TEV cleavage site. The DNA sequence was optimized for expression in *E. coli*.

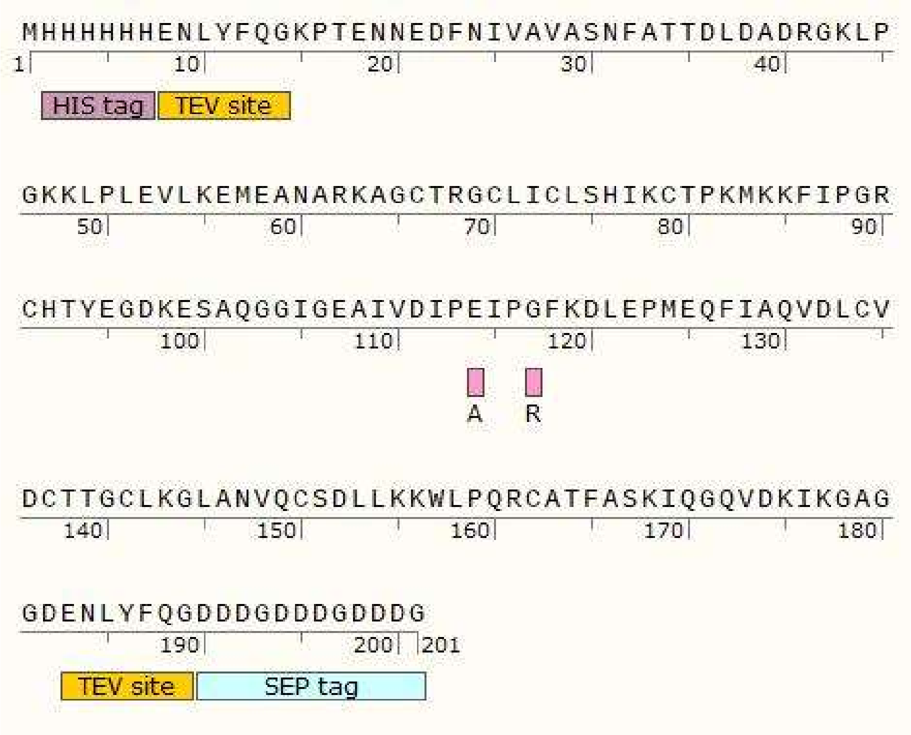


**Supplementary figure 1:** Sequence of GLuc as used in this project. The two spots where our sequence differs from the wild type sequence are shown in pink. (E100A and G103R).

The expression vector we used was a pET-21c expression vector, which featured an Ampicillin resistance gene.

We first transformed the BL21(DE3) *E. coli* strain with our new construct and later, after problems with the disulfide bond formation were observed, a BL21(DE3) containing the CyDisCo plasmid developed by the Ruddock group at Oulu university (Matos et al., 2014). **Supplementary figure 2** shows that this plasmid contains a polycistronic expression vector, pMJS205, for co-expression of the sulfhydryl oxidase, Erv1p, and the human Protein Disulfide-Isomerase (hPDI). It also contained the cat-gene for resistance towards chloramphenicol (Chlor).

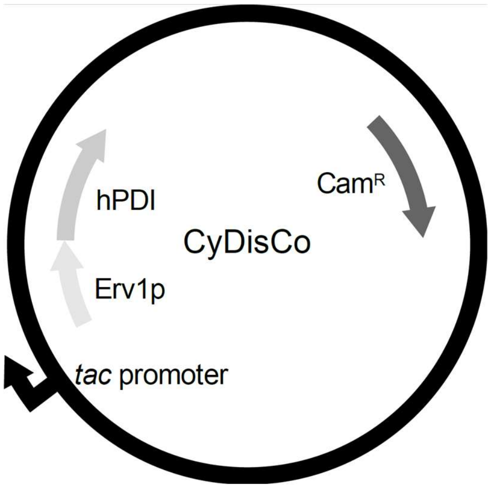


**Supplementary figure 2:** CyDisCo plasmid, pMJS205 (Gaciarz et al., 2016)

Both cell lines were transformed using the heat-shock method. Glycerol stocks of the resulting cells were kept at 80 °C until needed for expression.

#### Expression and purification

To express GLuc, 10 ml liquid LB precultures starting from a single colony were made the day before expression and grown overnight at 37 °C. The day of expression 1 liter freshly mixed AB-LB medium at room temperature was inoculated with a single preculture. The culture was grown shaking at 37 °C to an optical density of between 0.6 AU and 0.7 AU, after which expression was induced by adding IPTG to a concentration of 0.2 mM. The culture was then left shaking at 37 °C overnight. The next morning, the cells were spun down at 8000 g, for 15 min at 4 °C. The pellet was kept at -20 °C until purification. All media contained 100 µg $ml^{-1}$ ampicillin and the media used for expression with the CyDisCo system contained an additional 50 µg $ml^{-1}$ chloramphenicol.

The cell pellet was resuspended in 25 ml lysis buffer and sonicated 8 x 30 s at 50 % amplitude while on ice, repeated as necessary until homogeneity, using a UP200S ultrasonic processor (Hielscer). The lysate was then spun at 15 000 g and 4 °C for 15 min. The pellet was discarded. DNA in the supernatant was removed by adding streptomycin to a concentration of 1 % and stirring on ice for 30 min to allow a precipitate to form. The precipitate was then removed by spinning for 30 min at 15 000 g. and 4 °C. The supernatant was kept for immobilized metal affinity chromatography (IMAC), which was performed on an ÄKTA FPLC equipped with a column packed with 2 ml Thermo Scientific His-Pur Ni-NTA Superflow beads, which was equilibrated with wash-buffer beforehand. The extract was loaded onto the column in wash buffer at a flow speed of 0.5 ml $min^{-1}$ and then washed with another 2 column volumes of wash buffer. The protein was eluted over a gradient with 600 mM imidazole. The eluent was collected in 2 ml fractions and the protein elution was followed by UV absorption and conductivity. The elution peak was typically observed around 25 % elution buffer.

After purification, the fractions containing protein were combined. Ammonium sulfate (AMS) was added to a concentration of 50 % to precipitate impurities. After equilibrating on ice for 30 min the precipitate was spun down at 14 000 g, 4 °C for 15 min. The concentration AMS of the supernatant was then increased to 75 % to precipitate the protein. After another 30 min on ice the protein was spun down again at 14 000 g, 4 °C for 15 min and the supernatant was removed. The pellet was dissolved in 500-1000 µl storage buffer and dialyzed against 2 liter 50 mM Tris pH 8.0 at 4 °C two times for at least 2 h each.

GLuc concentrations were estimated from absorbance at 280 nm, using a theoretical extinction coefficient of 10595 $M^{-1}cm^{-1}$ calculated with the Protparam-tool on the ExPaSy server (Gasteiger et al., 2005).

#### Comparative expression with and without CyDisCo system

In the comparison between expression with and without the CyDisCo system an adjusted protocol was used. 200 ml cultures of both strains were grown in AB-LB medium. The used antibiotic concentrations were 50 µg $ml^{-1}$ for Ampicillin and 50 µg $ml^{-1}$ for Chlor. Cultures were incubated at 37 °C until they had reached an OD of between 0.8 AU and 1.0 AU. Then for 4 more hours at 37 °C, followed by incubation at 25 °C overnight.

Sample purification followed the description above in "Expression and purification", but IMAC was performed on a gravity column instead of on the ÄKTA FPLC. The protein was eluted with 300 mM imidazole, after which AMS purification was performed as in section 2.3. 250 µl samples were taken from both the 50 % and 75 % AMS suspensions. They were spun down and after separation of pellet and supernatant the pellets were dissolved in 250 µl TBS buffer pH 8.0. 27 µl of each sample was precipitated with TCA. 10 µl of each sample was loaded on a 14 % SDS gel. The gel was stained with coomassie stain.

#### Luminescence assays: general

All luminescence assays were performed in 50 mM TRIS buffer, pH 8, with 0.1 g $l^{-1}$ BSA to prevent adhesion of luciferase to the cuvette walls. Light emission at 480 nm was measured using a Perkin Elmer LS55 Luminescence spectrometer with 3 ml stirred cuvette, thermostated at 25 °C, with a slit width of 20 nm, delay of 1 ms, gate time of 20 ms, cycle time of 20 ms, flash count of 1 and a data interval of 0.1 s. Unless stated otherwise, 20 µl of luciferase solution in buffer was added to 1960 µl of assay buffer before starting the measurement. To catch the first part of the luminescence peak, we wanted to be able to inject coelenterazine without opening the lid of the spectrometer. Therefore, we custom-designed a black syringe holder which allowed injection through the lid, directly into the cuvette (**supplementary figure 3**). A syringe was filled with 20 µl of coelenterazine solution in isopropanol and placed in the syringe holder before starting the measurement. 10 s after the start of the measurement, the coelenterazine was injected into the reaction mixture, bringing the total volume to 2000 µl and the isopropanol concentration to 1 %. For consistency in CTZ concentration, all CTZ solutions were kept at -20 °C until right before the start of the assay. During the preparation of the injection, GLuc and CTZ were kept on ice.

Since it was found that isopropanol inhibits GLuc (**supplementary figure 6**), we always injected the same volume of CTZ solution, so the total isopropanol concentration in the assay was 1 %.

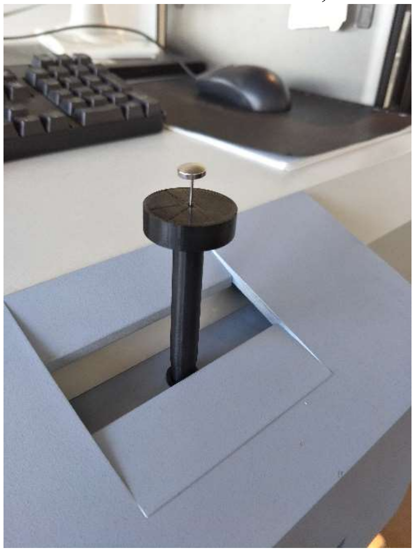

**Supplementary figure 3:** Syringe holder

#### NanoLuc expression and purification

5 ml liquid LB precultures, started from a single colony of BL21(DE3) cells containing plasmid pBB817 with a single copy of the full-length Oplophorus luciferase gene, were made a day before expression and grown overnight at 37 °C.
500ml AB-LB medium was inoculated from the ON culture and grown at 37 °C to an optical density 0.5. At this point, the culture was induced with 500µl 1M IPTG, to a final concentration of 1mM. The culture was left to express at 25 °C for 3 hours.
Cells were harvested by centrifugation and sonicated on ice at maximum intensity for 8 x 30s. The resulting lysate was centrifuged and the supernatant decanted.

The NanoLuc was purified from lysate by immobilized metal affinity chromatography (IMAC), which was performed on an ÄKTA FPLC equipped with a column packed with 2 ml Thermo Scientific His-Pur Ni-NTA Superflow beads, which was equilibrated with 50 mM $Na_2PO_4$, 150 mM NaCl, pH 8 beforehand. After washing with the same buffer, but including 20 mM imidazole, the protein was eluted in steps, with increasing concentrations of imidazole at each step, as shown below:

| Step | [Imidazole] |
|---|---|
| Equilibration | 20 mM |
| Elution 1 | 58.4 mM |
| Elution 2 | 116 mM |
| Elution 3 | 500 mM |

Elution steps 1 and 2 contained the majority of the NanoLuc, at high purity. Fractions were collected, dialyzed into 30 mM TRis, 1 mM EDTA, pH 7.6), and stored at -20C.

#### NanoLuc assay

Assays were performed as described above and in the results section. A NanoLuc standard curve was made by doing seven assays with a NanoLuc concentration of 0.2 nM and CTZ concentrations in the range of 13 nM to 860 nM. The average light signals between 10 s and 20 s were fitted to a Michaelis-Menten equation. The resulting fit had a $K_M$ of 829 nM and a $V_{max}$ of 1749 RLU (**supplementary figure 7**).
For each of the NanoLuc assays, NanoLuc was injected into the reaction mixture when the light signal had gone down to 10 % of the initial peak height. To calculate how much CTZ was left at the point of NanoLuc injection, the average height of the NanoLuc signal between 10 s and 20 s was compared to the standard curve fit. This number was then corrected for CTZ auto-degradation by dividing by $e^{-0.0003t}$, which would be the fraction of the substrate lost by auto-degradation. In this, *t* is the time between the start of the GLuc signal and the start of the NanoLuc signal and the factor 0.0003 is the decay rate of CTZ in buffer, measured in a separate experiment (**supplementary figure 8**). The number of turnovers per enzyme molecule was calculated by first subtracting the amount of leftover CTZ from the amount present at the start of the reaction and then dividing this number by the amount of GLuc present in the reaction.
For the calculation of the conversion factor between light output and number of cycles, the integrals were normalized for enzyme amount in nmol. Since the NanoLuc assays were measured at 460 nm instead of 480 nm, like all other assays, we had to adjust for this difference. In a separate experiment we determined that 1.25 times less light was detected at 460 nm than at 480 nm for GLuc. This was factored into the conversion factor.

#### Assays for enzyme and substrate concentration dependency

37 assays were performed, consisting of combinations of 8 different substrate concentrations ranging from 0.18 µM to 10 µM and 12 different enzyme concentrations ranging from 51 pM to 28.7 nM. Table 1 shows the combinations. High CTZ concentrations were matched with low GLuc concentrations and vice versa in order to stay below the detection limit of our equipment. Each assay was performed in triplo. GLuc and CTZ solutions were made as a dilution series and aliquoted. GLuc

solutions were stored at 4 °C and CTZ at -20 °C until use. All assays were recorded within 8 days, in random order.

*Table 1: Combinations of concentrations of CTZ and GLuc that were used to produce assays. Left column and top row show all used concentrations. Crosses indicate an assay was performed.*

| | CTZ (µM) | | | | | | | |
|---|---|---|---|---|---|---|---|---|
| | 0.178 | 0.316 | 0.562 | 1.000 | 1.778 | 3.162 | 5.623 | 10 |
| GLuc (nM) | | | | | | | | |
| 0.051 | | | | | | | X | X |
| 0.091 | | | | | | | X | X |
| 0.16 | | | | | X | X | X | X |
| 0.29 | | | | X | X | X | X | X |
| 0.51 | | | | X | X | X | X | X |
| 0.91 | | | | X | X | X | | |
| 1.62 | | X | X | X | | | | |
| 2.87 | X | X | X | | | | | |
| 5.11 | X | X | X | | | | | |
| 9.06 | X | X | X | | | | | |
| 16.2 | X | X | X | | | | | |
| 28.7 | X | | | | | | | |

#### Data analysis and curve fitting

The luminescence data was saved as a text file (ascii) and analyzed with an in-house Python-based interface. The first 10 s of the measurement (before start of the signal) were used to calculate the background. The average background signal was then subtracted from all data points. To find the start of the light signal, the difference between each data point and the average of the 10 data points (1 s) before it was calculated. If the value exceeded the average with more than a threshold value (0.3 by default), a signal was detected. If the default threshold resulted in the detection of signals where no actual signal was started, it was increased by the user until only the real signal was detected. For fitting purposes, the time at the start of each signal was set to 0 s. To fit signals to a model curve, they were first integrated. Then they were fitted to a model curve using the optimize.curve_fit function from the SciPy module for Python. An initial guess for the parameters was provided by the user, for the other settings the default values were used. All other fits were performed using the same SciPy function (optimize.curve_fit). Scripts are available on request.

Supplementary figures

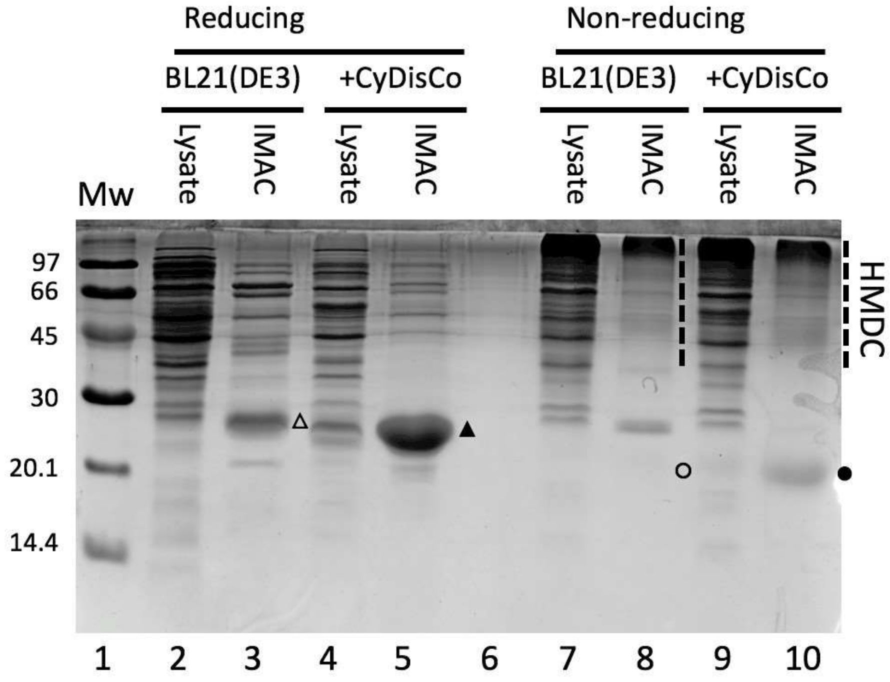


**Supplementary figure 4:** *SDS-PAGE under reducing and non-reducing conditions, comparing expression of SEP-tagged GLuc in BL21(DE3) in the absence and presence of the CyDisCo plasmid. Cell lysate and pooled IMAC eluate corresponding to equal volumes of original culture was run under reducing and non-reducing conditions as indicated. Under non-reducing conditions the correctly disulfide-linked material (closed circle; lane 10) moved faster than observed in the reduced samples (triangles, lanes 3 and 5) indicating a more compact disulfide-linked monomer. Lane 6 was left empty to avoid DTT cross-reaction. The gel was stained with Coomassie blue.*

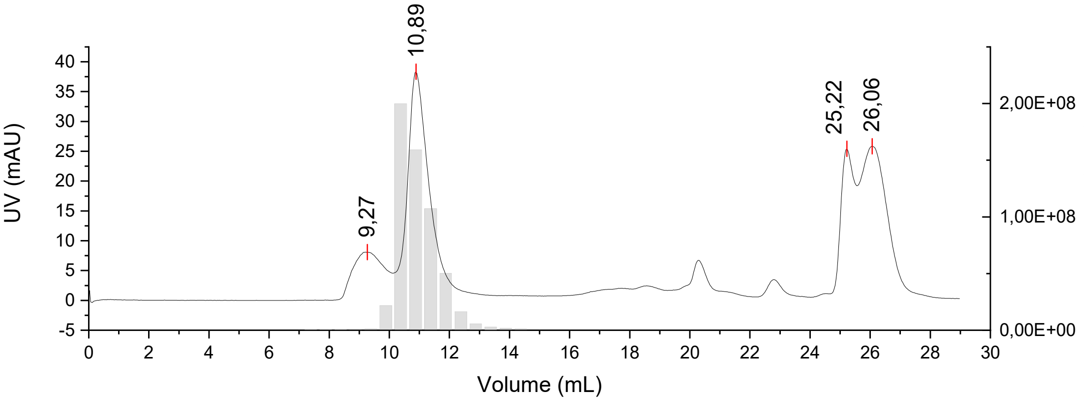


***Supplementary figure 5:*** *Size exclusion of Ni-NTA eluate. HMDC-GLuc elutes first, at 9.27 mL. GLuc elutes at 10.89 mL. Fraction of 0.5 mL between retention volume 7-15 mL were tested for activity. This shows that the HMDC-GLuc is inactive.*

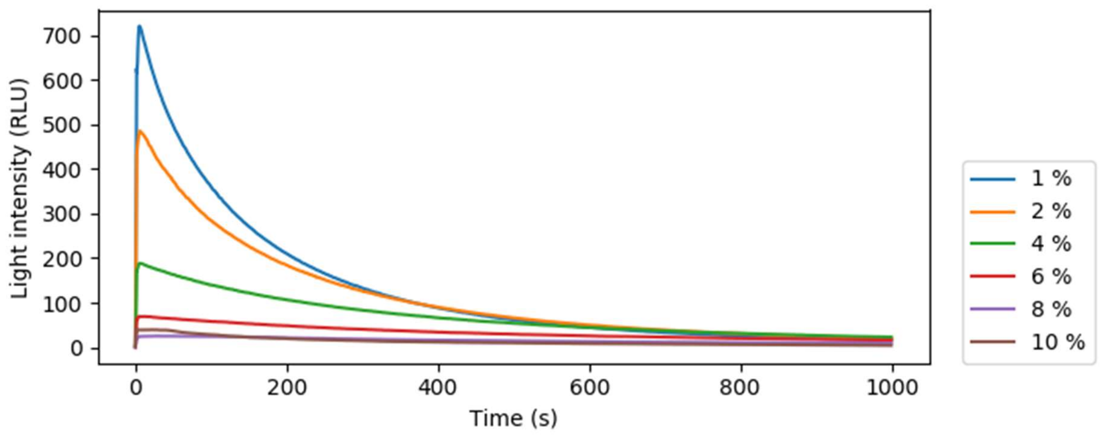


***Supplementary figure 6:*** *Isopropanol inhibits the GLuc reaction. The figure shows that the light output decreases with increasing isopropanol concentration.*

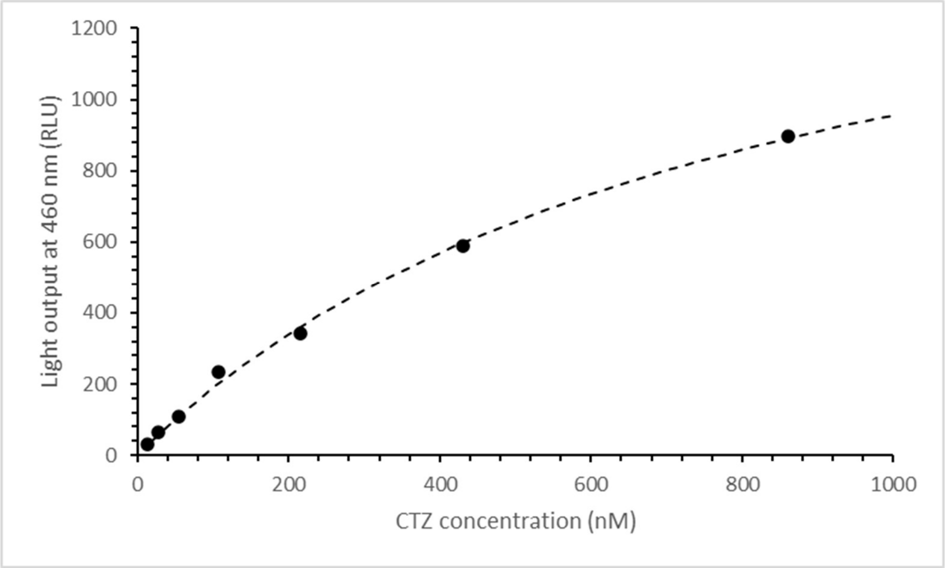


***Supplementary figure 7:*** *NanoLuc standard curve to relate the CTZ concentration in the cuvette to light output at 460 nm. The average light output between 10 s and 20 s after the start of the signal was used. The data was fitted to the Michaelis-Menten equation (dashed line), which was used as the standard curve. We found a* $V_{max}$ *of 1749 RLU and a* $K_M$ *of 829 nM.*

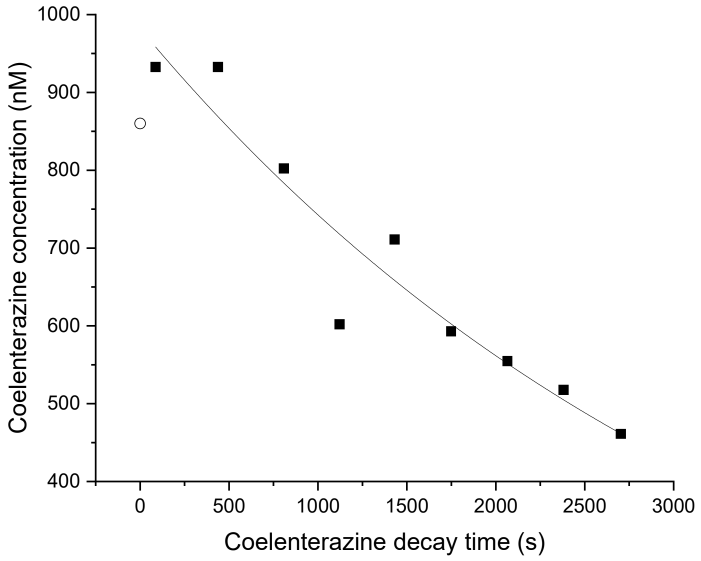


***Supplementary figure 8:*** *Coelenterazine (CTZ) auto-degrades in buffer. Here the decay rate was measured under standard assay conditions. Samples were taken out of the reaction mixture to measure light output at the times indicated in the graph. Light output was measured by injecting NanoLuc into 2 ml of sample. CTZ concentration was calculated from a NanoLuc standard curve (**supplementary figure** 7). The open circle indicates the CTZ concentration at the start, this was not measured by NanoLuc and is not included in the fit.*
*The data was fitted to the exponential decay curve* $ae^{-kx}$ *, resulting in an **a** of 981,95 ± 42,89 nM and a decay rate **k** of 2,79E-4 ± 3,34E-5* $s^{-1}$.

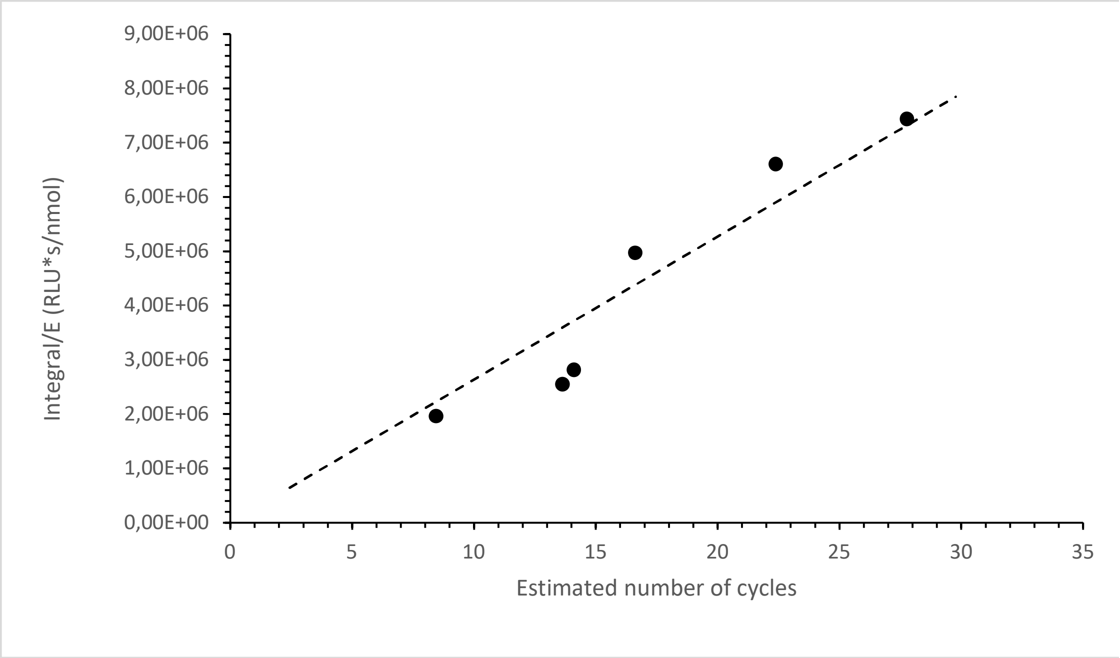


***Supplementary figure 9:*** *Cycles estimate*

*This figure demonstrates how we converted an integrated light signal to an estimate of the average number of cycles made by the enzyme before inactivating. The number of cycles in this graph was calculated from the NanoLuc assay, in which the amount of CTZ left after a GLuc reaction was measured by adding NanoLuc and recording the light signal.*
*The total integral a was normalized for the amount of enzyme present and corrected for light output at 480 nm instead of 460 nm. We measured in a separate experiment that the GLuc light signal is 1.25 times higher at 480 nm. Since the NanoLuc measurements were performed at 460 nm, we multiplied by 1.25 to obtain a more useful conversion factor.*
*For our equipment and under our assay conditions the number of reactions cycles relates to the amount of light with the following relationship:*

$$RC = \frac{Int}{n_{GLuc} * F}$$

*In which RC is the average number of reaction cycles per enzyme molecule; Int is the integral at 480 nm in RLU·s;* $n_{GLuc}$ *is the amount of GLuc partaking in the reaction in nmol and F is a device specific conversion factor, in our case* $2.64 \cdot 10^{-5}$ *RLU·s/nmol.*

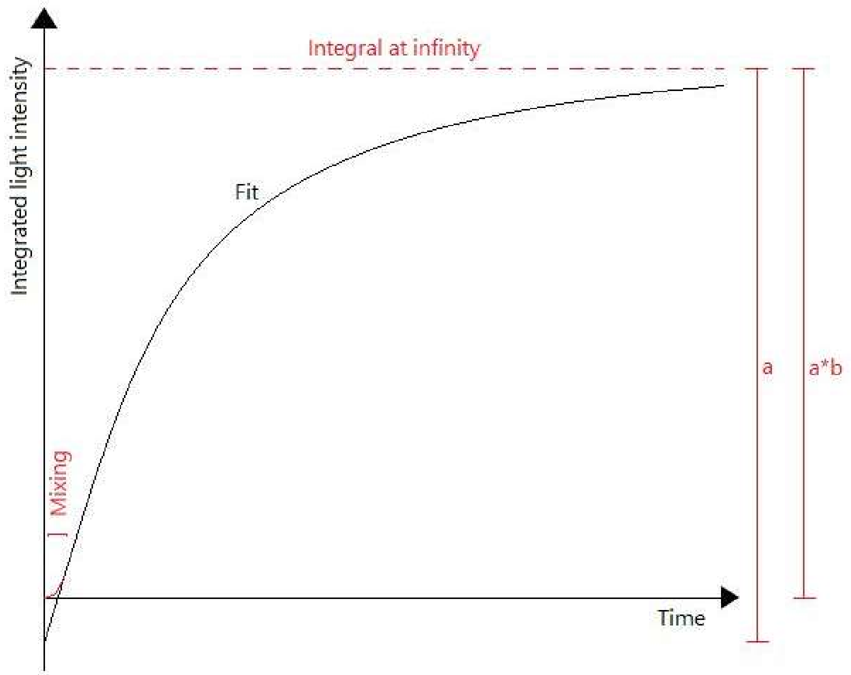


***Supplementary figure 10:*** *Demonstration of parameters a and b in exponential fits. Because there is a little bit of mixing time when CTZ is first injected, the fit does not exactly cross (0, 0). If it had, the integral would have been slightly higher or lower and that is why a takes this offset into account. To make the fit still end up at the same value as the signal at the end, parameter b is added, which takes care of this correction.*

a)

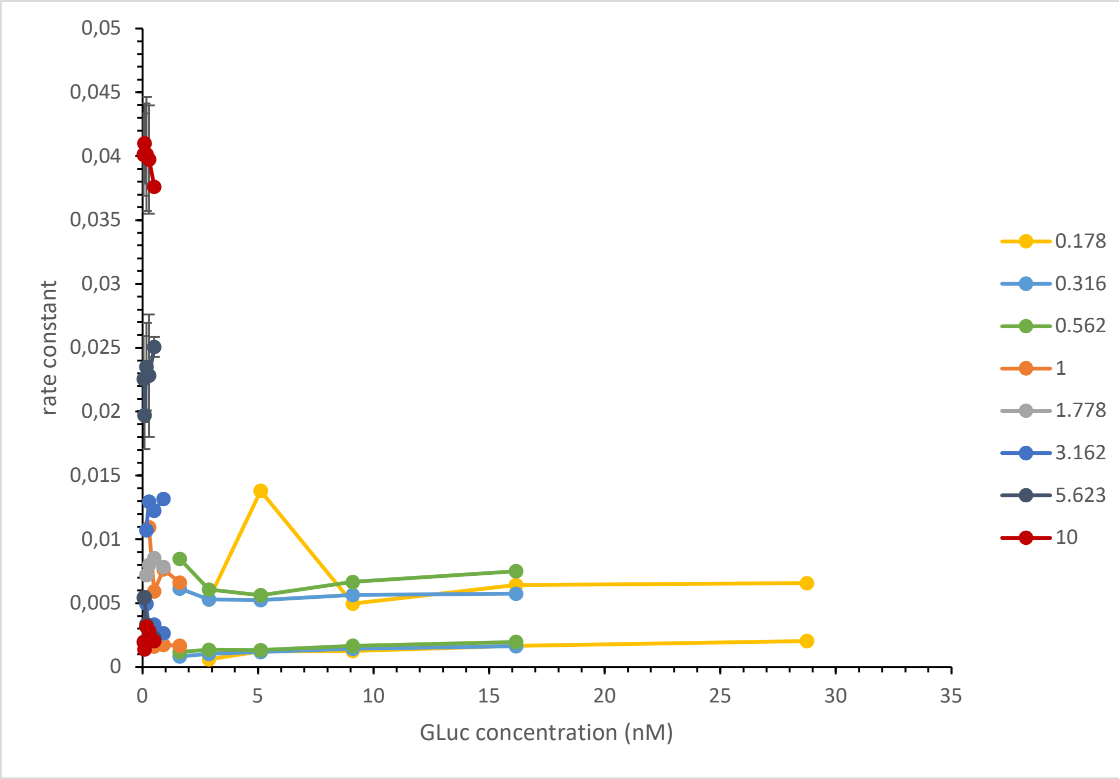

0,05
0,045
0,04
0,035
0,03
0,025
0,02
0,015
0,01
0,005
0
rate constant
0
5
10
15
20
25
30
35
GLuc concentration (nM)
0.178
0.316
0.562
1
1.778
3.162
5.623
10


b)

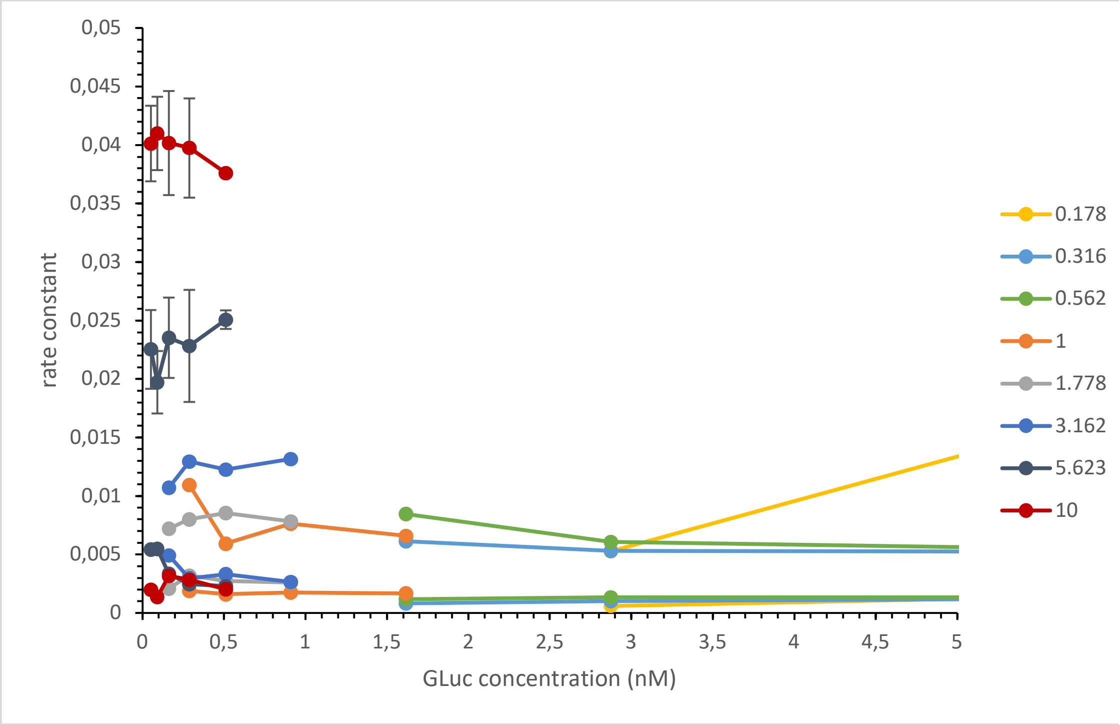

0,05
0,045
0,04
0,035
0,03
0,025
0,02
0,015
0,01
0,005
0
rate constant
0
0,5
1
1,5
2
2,5
3
3,5
4
4,5
5
GLuc concentration (nM)
0.178
0.316
0.562
1
1.778
3.162
5.623
10

***Supplementary figure 11:*** *Rate constants k1 and k2 plotted over the enzyme concentration. Full enzyme concentration range (a) and zoomed in on the lower enzyme concentrations (b). In both subfigures, the CTZ concentration is shown in different colors, with the rate constants k1 and k2 in the same color for the same substrate concentration. Each point was an average of three measurements. Error bars depicting standard deviations are shown for two series as a representative example. Both rate constants were constant with GLuc concentration.*

a)

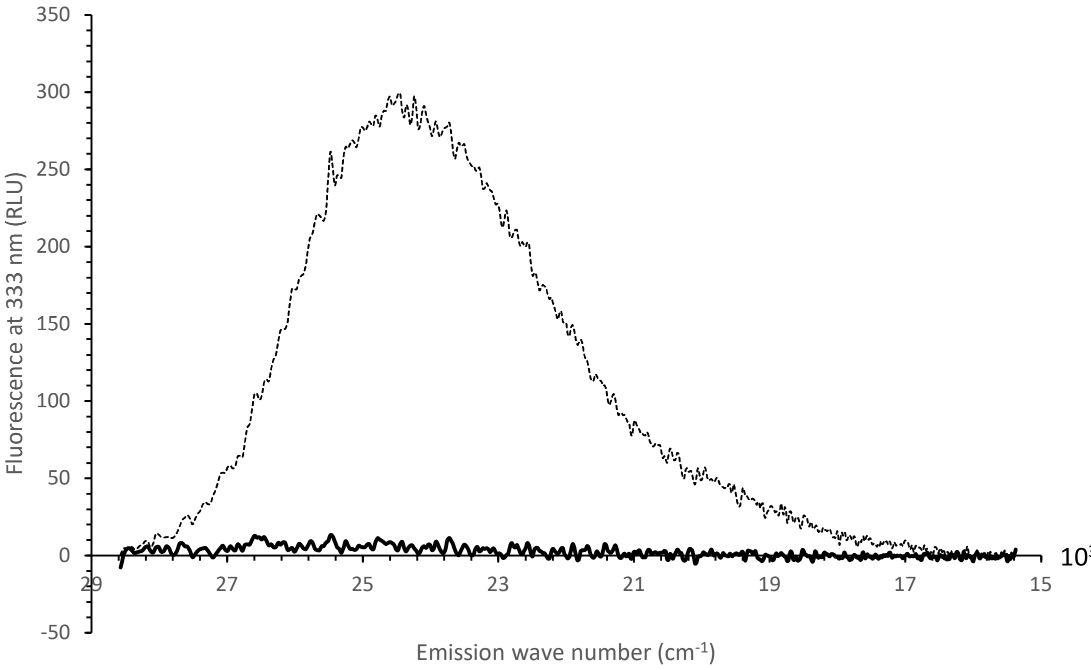


b)

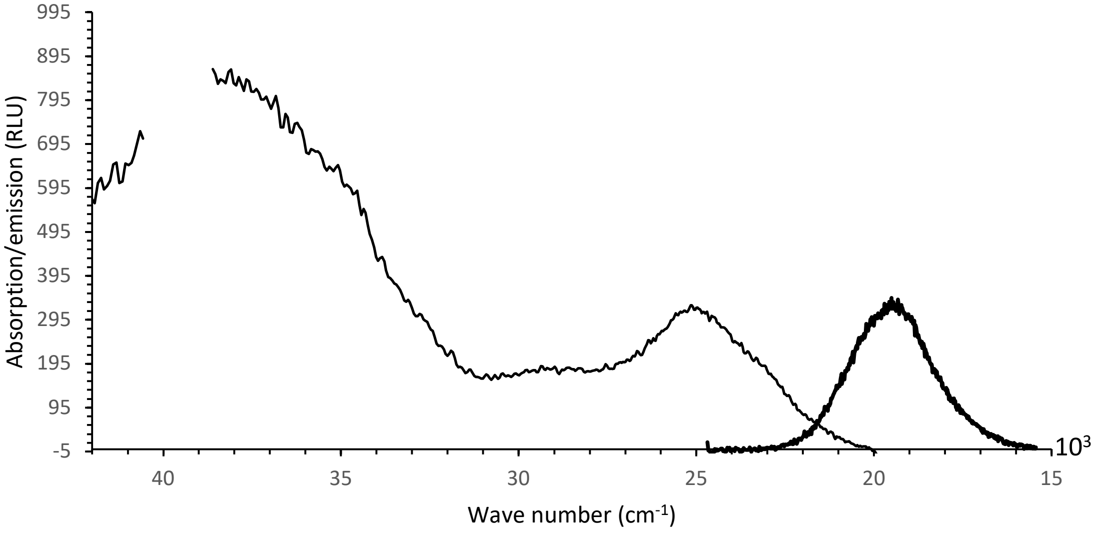

c)

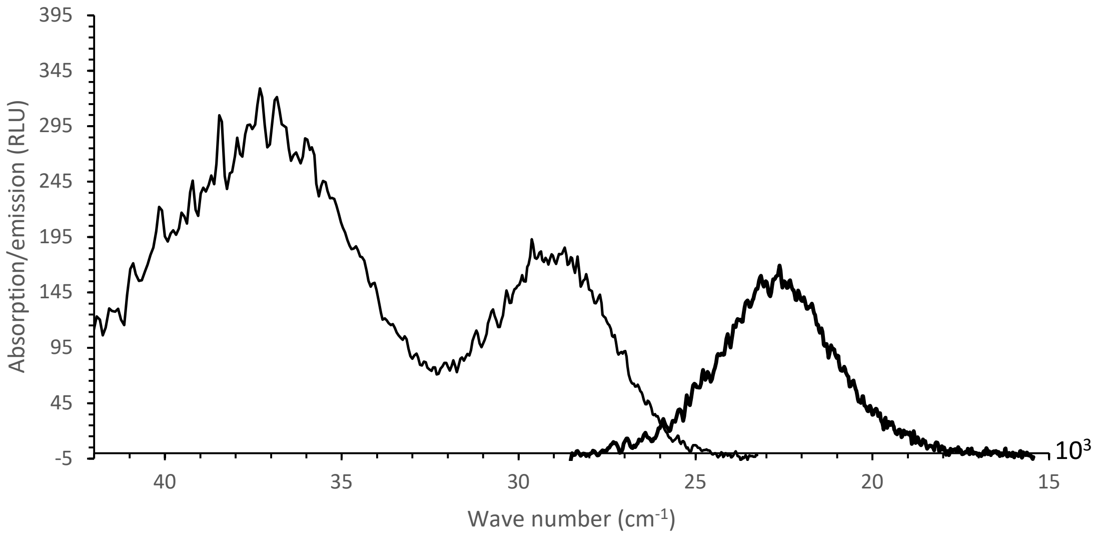


***Supplementary figure 12:*** *a) Emission spectra of active (bold) and inactivated (dashed) GLuc, excited at 333 nm/30.0·$10^3$ $cm^{-1}$. b) Excitation (emission fixed at 510 nm/19.6·$10^3$ $cm^{-1}$) and emission (excitation fixed at 395 nm/25.3·$10^3$ $cm^{-1}$) of fresh CTZ in aqueous buffer at pH 8. Data between 38.53·$10^3$ $cm^{-1}$and 40.5·$10^3$ $cm^{-1}$ was removed, because the solvent absorbed strongly here. c) Excitation (emission fixed at 440 nm/22.7·$10^3$ $cm^{-1}$) and emission (excitation fixed at 333 nm/30.0·$10^3$ $cm^{-1}$) of day-old CTZ in buffer at pH 8.*
